# Spatial organization and dynamics of RNase E and ribosomes in *Caulobacter crescentus*

**DOI:** 10.1101/228122

**Authors:** Camille A. Bayas, Jiarui Wang, Marissa K. Lee, Jared M. Schrader, Lucy Shapiro, W.E. Moerner

## Abstract

We report the dynamic spatial organization of *Caulobacter crescentus* RNase E (RNA degradosome) and ribosomal protein L1 (ribosome) using 3D single particle tracking and super-resolution microscopy. RNase E formed clusters along the central axis of the cell, while weak clusters of ribosomal protein L1 were deployed throughout the cytoplasm. These results contrast with RNase E and ribosome distribution in *E. coli*, where RNase E co-localizes with the cytoplasmic membrane and ribosomes accumulate in polar nucleoid-free zones. For both RNase E and ribosomes in *Caulobacter*, we observed a decrease in confinement and clustering upon transcription inhibition and subsequent depletion of nascent RNA, suggesting that RNA substrate availability for processing, degradation, and translation facilitates confinement and clustering. Moreover, RNase E cluster positions correlate with the subcellular location of chromosomal loci of two highly transcribed ribosomal RNA genes, suggesting that RNase E’s function in ribosomal RNA processing occurs at the site of rRNA synthesis. Thus, components of the RNA degradosome and ribosome assembly are spatiotemporally organized in *Caulobacter*, with chromosomal readout serving as the template for this organization.

---

In bacteria, the RNA degradosome mediates the majority of messenger RNA (mRNA) turnover and ribosomal RNA (rRNA) and transfer RNA (tRNA) maturation.^1^ The RNA degradosome assembles on the C-terminal scaffold region of the RNase E endoribonuclease.^2^ RNase E in *Escherichia coli (E. coli)* and RNase Y, the RNase E homolog in *Bacillus subtilis (B. subtilis),* both associate with the cell membrane through a membrane-binding helix.^3-6^ One proposed rationale behind the observed membrane association is to physically separate transcription within the nucleoid from RNA degradation, thus providing an inherent time delay between transcript synthesis and the onset of transcript decay, avoiding a futile cycle. *Caulobacter crescentus (Caulobacter)* is an alpha-proteobacterium widely studied as a model system for asymmetric cell division. Unlike *E. coli* or *B. subtilis, Caulobacter* contains a pole-tethered chromosome that fills the cytoplasm.^7-9^ Surprisingly, *Caulobacter* RNase E does not have a membrane targeting sequence and is not membrane-associated.^2, 10^ Furthermore, diffraction-limited (DL) images of RNase E exhibited a patchy localization throughout the cell, with RNase E directly or indirectly associating with DNA.^10^

Fluorescently labeled ribosomal proteins S2 and L1, proxies for ribosomes in *E. coli* and *B. subtilis*, respectively, were found enriched at the cell poles, spatially excluded from the nucleoid on the hundreds of nanometer scale.^11, 12^ Similar fluorescence imaging studies in *Caulobacter* revealed no apparent separation of ribosomes and the nucleoid, though did suggest that ribosome diffusion throughout the cell was spatially confined.^10^ However, because the *Caulobacter* cell is only ~500 nm by ~3.5 μm in size, the diffraction-limited resolution of conventional fluorescence microscopy would obscure any organization or motion of the components of the degradosome or the ribosome that are on length scales of less than ~200 nm.

We used a combination of live cell single particle tracking (SPT) and fixed cell superresolution (SR) microscopy to study the dynamics and spatial distribution of eYFP-labeled^13^ RNase E and ribosomal protein L1 in *Caulobacter* on sub-diffraction limit length scales. SPT and SR provide improved resolution down to ~20-50 nm using fluorescent proteins (FPs).^14, 15^ Moreover, both SPT and SR are single-molecule (SM) methods, allowing us to investigate the heterogeneity in protein behavior and distribution across many different cells. To avoid artifacts from projecting SM localizations onto two dimensions, we have used the double-helix point spread function (DH-PSF) which encodes axial information in each single-molecule image.^16^

Using these methods, we observed that the number of RNase E clusters changed as a function of the cell cycle, while the size of the clusters remained constant. However, ribosomes formed only weak clusters which were deployed throughout the cell, as we found to be the case in another alpha bacterium, *Sinorhizobium meliloti (S. meliloti).* For both RNase E and ribosomes, we observed a decrease in confinement and clustering when transcription was inhibited, likely due to depletion of RNA substrates for degradation, processing, and translation. We found that the localization of RNase E clusters correlates with the subcellular chromosomal position of the highly expressed rRNA genes that are positioned at two distinct sites on the chromosome, consistent with processing occurring at the site of rRNA synthesis. Thus, co-transcriptional activities exhibit dynamic subcellular organization that is dependent on the site of chromosomal readout.

## Results

*Diffusion of RNase E and ribosomal L1 proteins is influenced by transcriptional activity*

### RNase E

The endoribonuclease RNase E forms the core and scaffold of the *Caulobacter* RNA degradosome. Other components include the exoribonuclease polynucleotide phosphorylase, a DEAD-box RNA helicase RhlB, the Krebs cycle enzyme aconitase, and the exoribonuclease RNase D.^2, 17^ To study the motional dynamics of RNase E, we performed 3D SPT in live *Caulobacter* cells using a strain expressing RNase E-eYFP as the only copy of RNase E. We performed an initial photobleaching and waited for stochastic recovery and blinking of eYFP SMs^18^ (Fig. 1a, left). We then obtained 3D trajectories of individual molecules by joining DH-PSF localizations over consecutive frames (Methods, Fig. 1a, right). In contrast to *E. coli* in which RNase E diffuses on the membrane,^3^ RNase E molecules in *Caulobacter* were found throughout the cytoplasm of the cell, diffusing limited distances (Fig. 1b and 1d).

**Figure 1:**
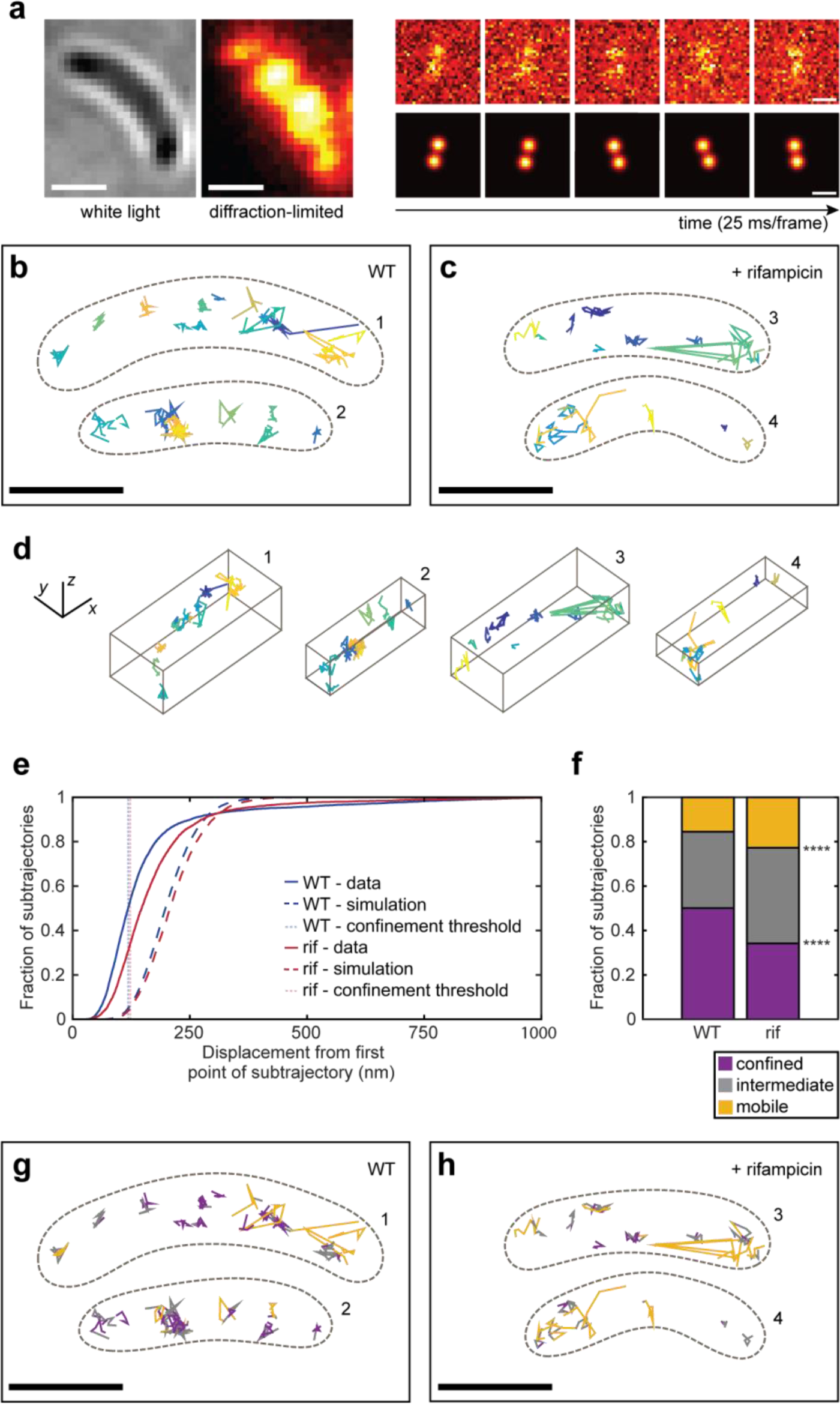
SPT of RNase E molecules measures decreased confinement when transcription is inhibited by rifampicin. **a**, left, Representative WL and DL images of a live *Caulobacter* cell expressing RNase E-eYFP. The DL image displayed was taken during the initial bleachdown of eYFP. **a**, right, Examples images of SMs in subsequent frames used to produce a SM trajectory. The DH-PSF experimental raw data (top row) matches well to the fit to two 2D Gaussians done through easy-DH-PSF (bottom row). The midpoint of the two lobes provides *xy* position, while the angle between the two lobes encodes axial position. **b**, Two representative WT cells with their RNase E trajectories. **c**, Two representative rif-treated cells and trajectories. Each trajectory is plotted as a different color. Only trajectories of at least 4 steps (5 frames or 125 ms) are shown and analyzed. The cell outline is indicated by the dashed brown line. **d**, 3D perspective views of the trajectories in b and c. **e**, CDF of displacements used for confinement analysis. Thresholds for classifying a subtrajectory as “confined” or “mobile” were set each for WT and rifampicin-treated cells, and were chosen to be the 5th percentile of the Brownian simulations for confined (pink and light blue dashed lines) and 50^th^ percentile for mobile. All other subtrajectories are defined as “intermediate”. **f**, Results from confinement analysis. g and h, Same cells as in b and c but with the subtrajectories color-coded according to their assigned confinement type. Analysis was performed on 2817 trajectories from 302 WT cells, 2709 trajectories from 351 rifampicin-treated cells. Scale bars are 500 nm in **a** and **d** and 1000 nm in **b, c, g,** and **h**.

To study the effect of RNA substrate availability on RNase E dynamics, we treated the cells with rifampicin (rif), an antibiotic that blocks transcription and effectively depletes cells of RNA.^19^ The ensemble diffusion coefficient for wild-type (WT) RNase E was extracted by analysis of the ensemble-averaged mean-squared displacement (MSD) (Methods, Supplementary Fig. S1a). The average diffusion coefficient for WT cells was found to be 0.0337 ± 0.0011 μm^2^/s (error is standard error of the mean (SEM) as determined from 500 bootstrapped samples of individual tracks). The diffusion coefficient increased upon treatment with rif (0.0407 ± 0.0004 μm^2^/s) (Fig. 1c). There is a heterogeneity of diffusion coefficients across different trajectories (Supplementary Fig. S2) likely due to sampling a mixed population of cells in various developmental stages, stochastic gene expression between individual cells, and RNase E molecules not all acting on RNA (Supplementary Fig. S3 shows additional example cells).

As the change in the ensemble diffusion coefficient was not drastic, we analyzed individual trajectories of WT and rif-treated cells for their degree of confinement. Each trajectory was divided into sub-trajectories and the maximum displacement from the first point of each sub-trajectory was calculated^20, 21^ (Fig. 1e). Displacements were compared to simulated Brownian motion inside a cell volume using ensemble WT and rif diffusion coefficients (Supplementary Fig. S4) to control for the possibility that trajectories appear confined by chance or due to the cell shape. A threshold was set at the Brownian 5^th^ percentile, such that only 5% of the simulated Brownian subtrajectories are confined. Sub-trajectories with displacements less than this value were classified as confined. To include the possibility of an intermediate degree of confinement, another threshold was arbitrarily set at the Brownian 50^th^ percentile, and sub-trajectories with displacements greater than this value were classified as clearly mobile. All other sub-trajectories (between 5^th^ and 50^th^ percentile) were classified as intermediate. Upon rif treatment, the fraction of confined molecules decreased from 50.1% in WT sub-trajectories to 34.2%, while the fraction of mobile subtrajectories increased from 15.6% in WT to 22.8% (Fig. 1f-h).

Ensemble-averaged MSD’s for the confined and mobile WT sub-trajectories were analyzed (Supplementary Fig. S5) yielding drastically different D values of 0.0025 ± 0.0001 μm^2^/s (confined) and 0.1944 ± 0.0010 μm^2^/s (mobile). For rif-treated cells, the diffusion coefficient for confined sub-trajectories is 0.0060 ± 0.0001 μm^2^/s and for mobile sub-trajectories is 0.1270 ± 0.0072 μm^2^/s. The diffusion coefficient for WT confined sub-trajectories is consistent with a stationary particle, whereas the diffusion coefficient for rif confined sub-trajectories is larger than would be expected for a stationary particle given our localization precision (32/33/49 nm in *x/y/z).*

The difference in diffusion between WT and rif-treated cells is not an artifact due to a potential rif-induced chromosome structure disruption.^22^ We performed the same 3D SPT of RNase E molecules in a ΔHU2 background with significantly reduced nucleoid compaction.^23^ Results show similar confinement as that of WT cells (Supplementary Fig. S6), arguing against the possibility of a difference in chromosome packaging between WT and rif-treated cells contributing to the difference in RNase E diffusion.

### Ribosomes

To obtain information about ribosome dynamics and translation, we performed 3D SPT of ribosomes through imaging eYFP-labeled ribosomal protein L1, and found that L1 diffuses much more freely throughout the cytoplasm than RNase E (Fig. 2a, Fig. 2c, Supplementary Fig. S7 for additional examples). Multiple populations of diffusing molecules were present by visual inspection. We calculated the squared displacements of the trajectories using a time lag of 1 frame (25 ms), then computed the empirical cumulative distribution function (CDF) of these squared displacements. The empirical CDF was then fit to a standard two-population model (Methods) (Fig. 2d-e, Supplementary Fig. S8). We found easily observable diffusion: in WT cells, 89% of the population was found to have a diffusion coefficient of 0.039 ± 0.001 μm^2^/s (larger than would be expected for a stationary particle) (error is SEM as determined from 500 bootstrapped samples of individual squared displacements), while 11% of the population was found to have a much larger diffusion coefficient of 3.23 ± 0.12 μm^2^/s. The slower value is consistent with reported values of diffusion coefficients from SPT of *E. coli* ribosomes.^24, 25^ We hypothesize that the slow-diffusing population (89% of trajectories measured) are involved in active translation that slows down the translation machinery. The faster-diffusing population (11% of total trajectories) are likely free L1 subunits that have not been assembled into ribosomes.

**Figure 2:**
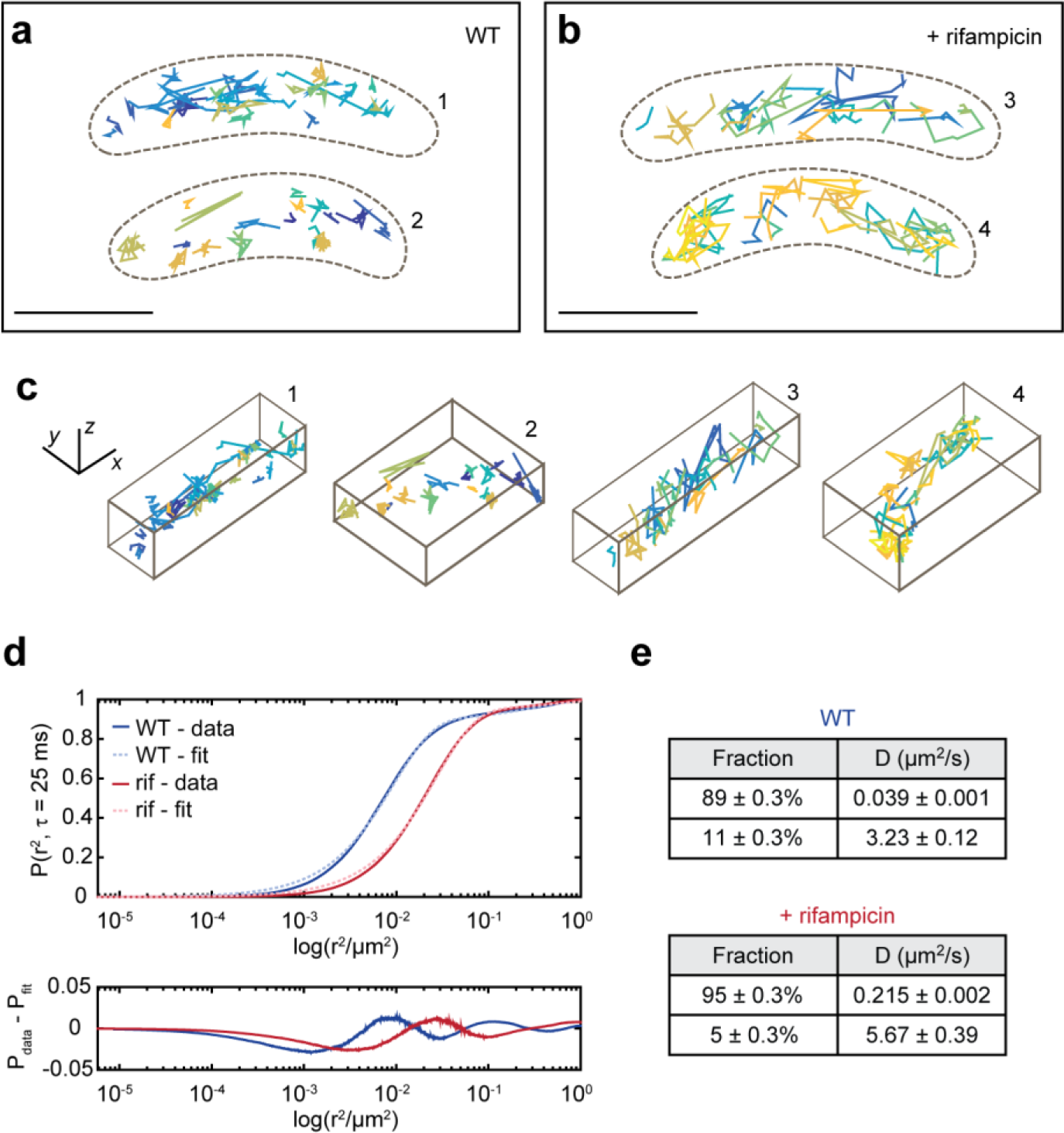
SPT of single ribosomal L1 molecules reveals faster diffusion in cells in which RNA has been depleted, as well as free- and ribosome-bound L1. **a**, Example WT cells and their trajectories. **b**, Example rif-treated cells and their trajectories. Each trajectory is plotted as a different color. Only trajectories of at least 4 steps (5 frames or 125 ms) are shown and analyzed. **c**, 3D perspective views of the trajectories in a and b. d, top, Semi-log plot of the CDF of squared displacements. **d**, bottom, Residuals of the fit to the CDF of squared displacements. **e**, Summary of the two diffusing populations in WT and rif-treated cells. Errors are standard deviations from 500 bootstrapped samples. Scale bars in a and b is 1000 nm and in c is 500 nm. Analysis was performed on 11249 trajectories from 298 WT cells and 12757 trajectories from 326 rifampicin-treated cells.

When cells are treated with rif (Fig. 2b-c), 95% of the population was found to have a diffusion coefficient of 0.215 ± 0.002 μm^2^/s, while 5% of the population was found to have a diffusion coefficient of 5.67 ± 0.39 μm^2^/s. Both populations in rif-treated cells have diffusion coefficients that are greater than observed for WT cells. The faster movement of the slow fraction in rif-treated cells may be attributed to ribosomes that have not yet bound RNA to process.

#### RNase E and ribosomal L1 proteins are differentially clustered in Caulobacter

As our SPT data have shown, RNase E and ribosomes both display dynamic behaviors in the cell. To study their spatial distribution without the confounding effect of motion, we turned to SR imaging of fixed cells. The imaging time required to obtain SR images with sufficient sampling is longer than the persistence time of individual mRNAs.^26, 27^ Accordingly, the cells were fixed to capture global RNase E or ribosome locations at a specific time window. Because the photophysical properties of eYFP (stochastic and reversible recovery from a long-lived dark state) can cause artifacts which mimic protein clustering,^28^ we removed localizations that were likely due to oversampling (Methods).

### RNase E

We first performed 3D two-color SR imaging of RNase E and the cell surface to confirm that *Caulobacter* RNase E is not sequestered to the cell membrane.^10^ In sharp contrast to RNase E sequestration to the inner membrane in *E. coli*,^4, 5^ RNase E in *Caulobacter* was observed to cluster along the central axis of the cell and nucleoid (Fig. 3a-d), consistent with previous diffraction-limited imaging studies showing the cytoplasmic localization of RNase E.^10^ RNase E formed distinct clusters in the cytoplasm (Fig. 3e, Supplementary video 1), although we observed various degrees of clustering across the population due to stochastic sampling as well as cell-to-cell variability in gene expression (Supplementary Fig. S10).

**Figure 3.**
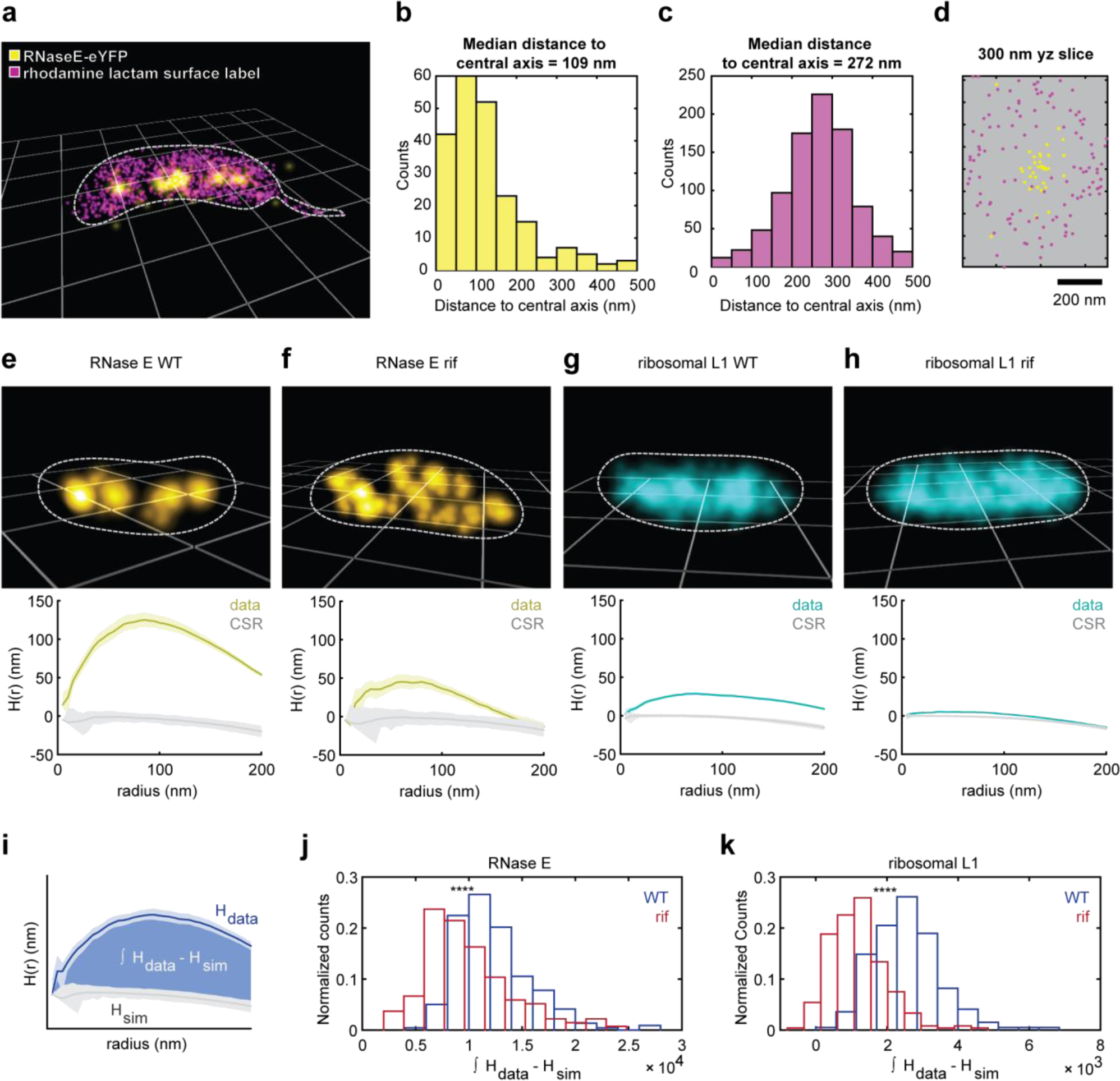
Spatial clustering of RNase E and ribosomes decreases when RNA is depleted. **a**, Example fixed cell expressing RNase E-eYFP (yellow) with its cell surface labeled with rhodamine spirolactam (magenta). **b**, Distribution of distances of RNase E molecules to the cell’s central axis. **c**, Distribution of distances of rhodamine lactam molecules to the cell’s central axis. **d**, A yz projection of a 300 nm slice perpendicular to the cell axis. **e**, SR reconstruction of a fixed WT *Caulobacter* cell expressing RNase E-eYFP, with calculation of the H(r) clustering metric (below) for the data as well as for CSR. **f**, SR reconstruction of a fixed rif-treated RNase E cell. **g**, SR reconstruction of a fixed WT *Caulobacter* cell expressing ribosomal protein L1-eYFP. **h**, SR reconstruction of a fixed rif-treated L1 cell. In the H function curves plotted in e-h, the lighter grey and lighter yellow (e-f)/lighter blue (g-h) show the 95% confidence intervals. In the reconstructions, each localization is plotted as a 3D Gaussian with a sigma equivalent to the average *xy* localization precision of the localizations in each cell (29/30 nm in *xy* for RNase E and 24/25 nm in *xy* for ribosomes). An average of 120 molecules and 720 molecules are plotted for RNase E and ribosomes, respectively. Cells are on 1 micron grids. **i**, The degree of clustering quantity plotted in f and g were calculated from the area between the data curve and CSR. **j**, Distributions for RNase E WT and rif-treated cells showing clustering relative to CSR. **k**, Distributions for L1 WT and rif-treated cells showing clustering relative to CSR. Analysis was performed on 218 WT and 135 rif-treated RNase E cells, and on 195 WT and 239 rif-treated ribosome cells.

RNase E in fixed *Caulobacter* cells was more clustered than would be expected from complete spatial randomness, indicating that the observed clustering is not due to chance or the cell shape^29^ (Methods). Positive deviations from the gray curve (Fig. 3e, bottom row) indicate clustering. The clustering was heterogeneous across the mixed cell population, likely arising from both intrinsic and extrinsic noise^26, 30^ (Fig. 3j). When transcription was inhibited by rif, RNase E clusters dissipated and the observed clustering shifted down towards complete spatial randomness (Fig. 3f, Supplementary video 2). To compare WT and rif-treated cells, we calculated a single clustering metric from the integral between the two curves (Fig. 3i) for each cell that captures the difference between the observed clustering in the data and the artefactual small clustering in the complete spatial randomness simulations arising from the cell shape (Methods) (Fig. 3j). The rif-treated values were significantly lower than those for the WT cells, which captures the decreased clustering of molecules in rif-treated cells (p < 0.0001).

### Ribosomes

We acquired SR images from fixed *Caulobacter* cells expressing ribosomal protein L1-eYFP (Fig. 3g, Supplementary video 3, Supplementary Fig. S11 shows additional example cells). Ribosomes were observed throughout the cytoplasm, consistent with diffraction-limited imaging^10^ and cryo-electron microscopy.^31^ This differs from the ribosome/nucleoid segregation observed in *E. coli* and *B. subtilis* ^24^ Ribosomes in fixed *Caulobacter* cells appear weakly clustered, yet slightly more clustered than what would be expected from complete spatial randomness (Fig. 3g, bottom row). Performing the same clustering analysis as we did with RNase E, we observed a heterogeneous distribution of clustering between different cells (Fig. 3k). Nevertheless, similar to RNase E, ribosome clustering exhibited a clear relationship with transcription, as treating the cells with rif significantly shifted the observed clustering towards complete spatial randomness (Fig. 3h and 3k, Supplementary video 4) (p < 0.0001).

To investigate whether cytoplasmic ribosomal distribution is unique to *Caulobacter,* we imaged ribosomal L1-eYFP in another alpha-proteobacterium, *S. meliloti*, together with the SYTOX orange DNA intercalating dye using epifluorescence microscopy. As a control, we also imaged an *E. coli* strain containing ribosomal protein S2-eYFP.^24^ Similar to *Caulobacter,* ribosomes in *S. meliloti* occupied the entire cell volume and co-localized with the nucleoid, whereas the gamma-proteobacterium *E. coli* maintained segregation of the DNA and polar nucleoid-excluded ribosomes (Supplementary Fig. S12). Notably, *S. meliloti* ribosomes contained a slightly more patchy localization pattern than *Caulobacter,* but generally showed a similar interpenetrating nucleoid-ribosome configuration similar to that observed with *Caulobacter.*

#### Number of clusters of RNase E changes as a function of the cell cycle

Ribosome profiling^32^ and Western blot analysis^2^ have shown that RNase E levels in *Caulobacter* vary as a function of the cell cycle, with maximum levels at the swarmer to stalked transition and just prior to cell division. To determine if RNase E clusters fluctuate as a function of the cell cycle, we performed two-color 3D imaging of RNase E-eYFP and PAmCherry-PopZ (to mark the cell poles). We imaged synchronized cells (Methods) to capture snapshots of the RNase E clusters at distinct time points along the cell cycle (Fig. 4a-e). Our 3D reconstructions show RNase E clusters along the cell axis throughout the cell cycle and PopZ foci at the cell poles.

**Figure 4:**
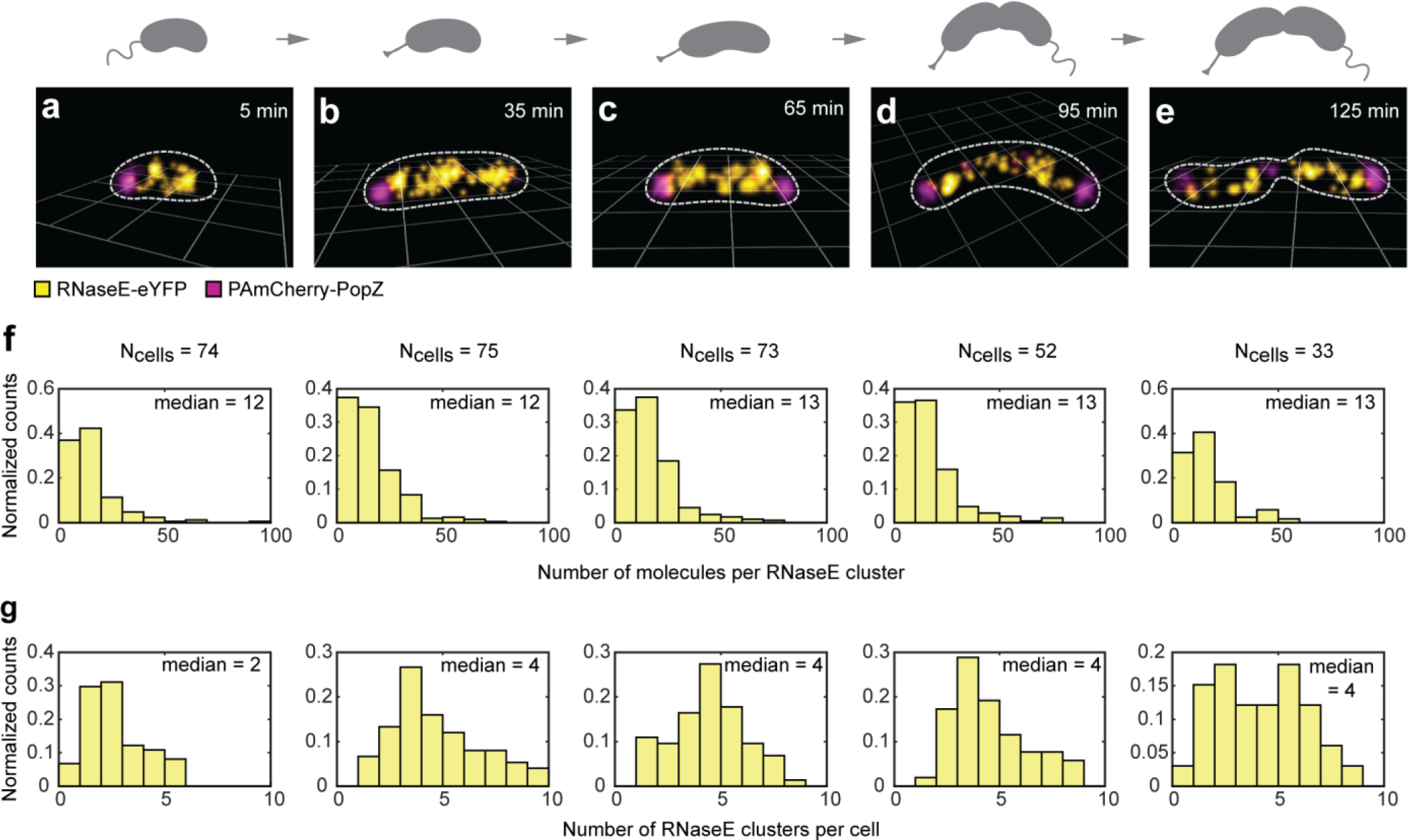
Cell-cycle dependent spatial distribution of RNase E-eYFP in fixed *Caulobacter* cells. **a-e**, SR reconstructions of fixed *Caulobacter* cells expressing RNase E-eYFP and PAmCherry-PopZ at (**a**) 5 minutes, (**b**) 35 minutes, (**c**) 65 minutes, (**d**) 95 minutes, and (**e**) 125 minutes after synchrony. The total cell cycle is 150 min. Cells are on 1 micron grids. **f**, Distributions of the number of molecules detected in each RNase E cluster throughout the cell cycle. **g**, Distributions of the number of RNase E clusters per cell throughout the cell cycle. We find a constant number of molecules in each cluster for all cell stages, but a doubling in the number of clusters per cell during the swarmer to stalk transition.

We then analyzed the sizes of RNase E clusters using the density-based algorithm DBSCAN (Methods) (Fig. 4f). There was no change in the median number of molecules detected in each RNase E cluster as cells progressed through the cell cycle. However, we did find an increase in the number of RNase E clusters as the swarmer cells differentiated into stalked cells (Fig. 4g). During the swarmer to stalked cell transition, the median number of clusters in the cell doubled from 2 clusters to 4 clusters in keeping with the previous observation that the abundance of RNase E varies during the cell cycle with a maximum at the swarmer to stalked cell differentiation.^2^ The histogram for the final 125-min population (Fig. 4g, right) shows the existence of two populations, since this cell type has both swarmer and stalked cells. The increase in the number of clusters in the cell during the swarmer to stalked transition is consistent with biochemical data showing increased overall RNA abundance during this time in the cell cycle.^32^ This increase in clusters may also be attributed to the initiation of DNA replication that occurs during this time period leading to an increase in overall transcription.

#### RNase E clusters co-localize with highly-transcribed genes

Our observation that the number of RNase E clusters decreased upon a decrease in RNA substrate availability after treatment with rif, suggested that RNase E clusters might form near regions of high transcriptional activity. Chromosomal loci along *Caulobacter’s* single circular chromosome occupy specific subcellular locations.^33^ Using this distinct organization of the chromosome, we were able to ask if RNase E clusters correspond to DNA loci that are highly transcribed. The genes encoding ribosomal RNA are the most highly transcribed loci, comprising 95-97% of the prokaryotic cell’s total RNA mass.^34^ There are two near-identical copies of rDNA in *Caulobacter* (Fig. 5a) that occupy spatially distinct sites on the chromosome. Each of these loci encode 16S, 23S, and 5S rRNA, as well as two tRNAs. These RNAs are transcribed as a single transcript that is subsequently processed by RNases, including RNase E, to generate the mature RNA species.^1, 2, 35^

**Figure 5:**
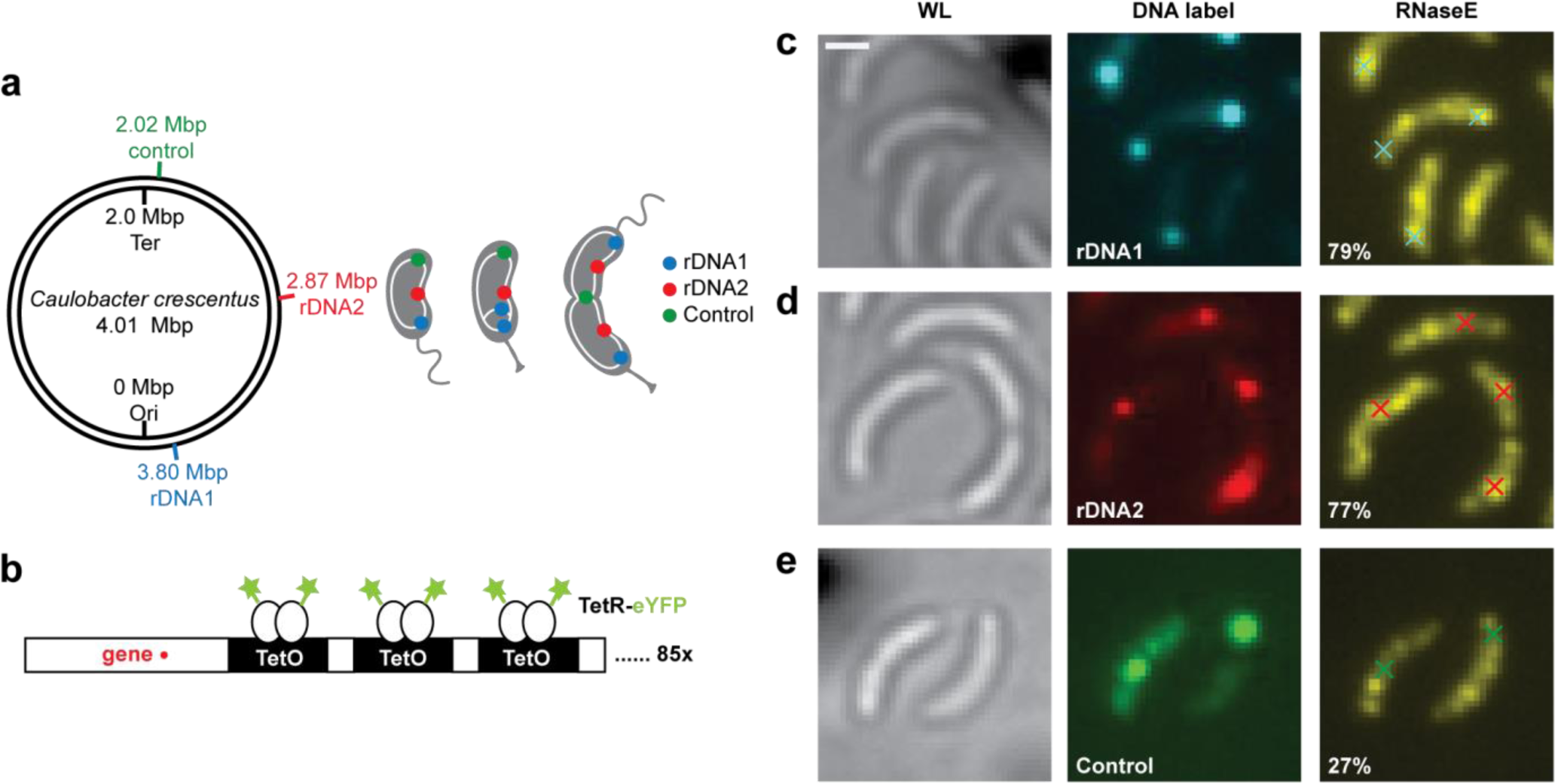
Colocalization of rRNA loci and CCNA01879 loci with RNase E. **a**, A cartoon of the labeled DNA loci. Approximate locations were calculated using the *Cauloabcter* swarmer 1D chromosome map as described previously.^33^ **b**, A schematic of the DNA-labeling scheme. c, left, A WL image of live *Caulobacter* cells expressing RNase E-PAmCherry and tetR-eYFP/85x-tetO on rDNA1 locus. **c**, middle, A DL image of the tetR-eYFP labeling the rDNA1 locus. c, right, A DL image of RNase E-PAmCherry. 79.2% of rDNA1 locus (cyan) colocalize with an RNase E cluster (yellow). **d**, *Caulobacter* cells with rDNA2 labeled. 76.9% of rDNA2 locus (red) colocalize with an RNase E cluster (yellow). **e**, *Caulobacter* cell with CCNA01879 labeled. 27.1% of the control CCNA01879 locus (green) colocalize with an RNase E cluster. Scale bars are 500 nm. Analysis was performed on 506 rDNA1 cells, 664 rDNA2 cells, and 321 control cells.

To determine if the observed RNase E clusters are positioned near the DNA loci of rRNAs, we labeled the two rDNA operons (rDNA1 and rDNA2) with a tetO/tetR labeling system (Fig. 5b), and visualized the positions of RNase E with respect to the two rRNA gene loci by diffraction-limited imaging. The positions of the rDNA gene loci were found by super-resolution imaging, and these positions were plotted against the RNase E diffraction-limited images (Fig. 5c-e) (Methods). We found that the two rDNA loci each co-localized with an RNase E cluster. Specifically, 79.2% of the first rDNA locus and 76.9% of the second rDNA locus co-localized with an RNase E cluster (additional examples are in Supplementary Fig. S13-S15). Thus, these results suggest that RNase E processing of ribosomal RNA transcripts occurs at the site of rRNA synthesis.

To confirm that these correlations are valid, we also imaged a lowly-transcribed gene, CCNA01879, that is in the bottom 0.5% of RNA-seq analysis^32^ (Fig. 5a). In contrast to rDNA, we found a lack of correlation of an RNase E cluster with this poorly transcribed locus. Only 27.1% of CCNA01879 loci co-localized with an RNase E cluster. There was still some co-localization, possibly because within any given cell, a few RNase E clusters are present which coincidentally can co-localize with our DNA labels. These results support the hypothesis that RNase E clusters are preferentially localized to highly-transcribed DNA loci where they immediately process RNA, most likely co-transcriptionally.

## Discussion

RNase E is the core of the RNA degradosome complex. It has been shown to be membrane-associated in *E. coli* and *B. subtilis*,^3-6^ but deployed throughout the nucleoid in *Caulobacter*.^10^ Previous studies of ribosome localization in *E. coli*^24, 25^ and *B. subtilis*^12^ revealed enrichment at the cell poles, where they are excluded from the nucleoid, while similar studies in *Caulobacter* revealed no apparent separation of ribosomes and the nucleoid.^10^ However, due to the small size of *Caulobacter,* the diffraction-limited resolution of standard fluorescence microscopy would obscure all sub-diffraction-limited motion or organization. Therefore, in this study we used 3D SPT and SR microscopy to probe and quantify the spatial organization of RNase E and ribosomes at high resolution, thereby providing insight into the dynamic behavior of these complexes.

Given that RNase E clusters were found in the cytoplasm in a pattern similar to that of the nucleoid, there is unlikely to be a gross physical separation of the RNA production and degradation machinery. We found the sub-cellular position of RNase E clusters correlated with sites of rRNA synthesis, indicating that cleavage occurs where RNA substrate is produced. This observation is consistent with evidence that the rRNA transcript is processed co-transcriptionally,^36^ and previous studies which have shown RNase E associating with DNA.^10^ The increased local concentration of RNase E clusters near highly-transcribed genes allow a large number of substrates to be quickly degraded or processed. RNase E clusters might reflect the formation of a large number of RNA degradosome complexes, which could facilitate efficient RNA cleavage. The size of these clusters did not change during cell cycle progression, suggesting that there might exist an optimum number of molecules per cluster. The observed confined sub-trajectories are likely actively processing RNAs in large RNA degradosome assemblies, possibly acting co-transcriptionally on transcripts still tethered to the chromosome. Mobile sub-trajectories could belong to RNase E molecules that have not yet bound an RNA substrate. Moreover, the decrease in the fraction of confined RNase E molecules and lower degree of confinement in cells in which RNA has been depleted highlights RNase E’s relationship with substrate availability.

We observed weakly clustered ribosomes throughout the cytoplasm, occupying the same regions as the nucleoid. This result is consistent with previous diffraction-limited imaging of ribosomes and the observation that transcription and translation occurs throughout the nucleoid,^10^ but contrasting with studies from *E. coli* and *B. subtilis.* For instance, ribosomes in *B. subtilis* are localized at regions distinct from the nucleoid due to actively-transcribed regions of the nucleoid being exposed to the cell periphery.^12, 37^ Upon substrate depletion through transcription inhibition, we observed a decrease in *Caulobacter* ribosome clustering and an increase in the diffusive population, suggesting a decreased fraction of ribosomes engaged in active translation. The weak clusters observed in WT cells as compared with rif-treated cells likely reflect a larger fraction of ribosomes engaged in translation. These weak clusters may act as translation hotspots as they can rapidly reinitiate translation on nearby mRNAs. We observed a sub-population of rapidly diffusing ribosomes in WT cells, which contrasts with prior FRAP experiments that observed essentially stationary ribosomes.^10, 38^ The diffraction-limited resolution of these previous measurements may have missed the fast-moving population of ribosomal protein L1, whereas SPT is able to gain higher resolution information.

Taken together, our results show that even without intracellular compartments, bacteria spatially organize RNA processing, degradation, and translation within the nucleoid similar to eukaryotic systems,^39^ and that this organization is governed by the transcriptional profile of the chromosome (Fig. 6).

**Figure 6:**
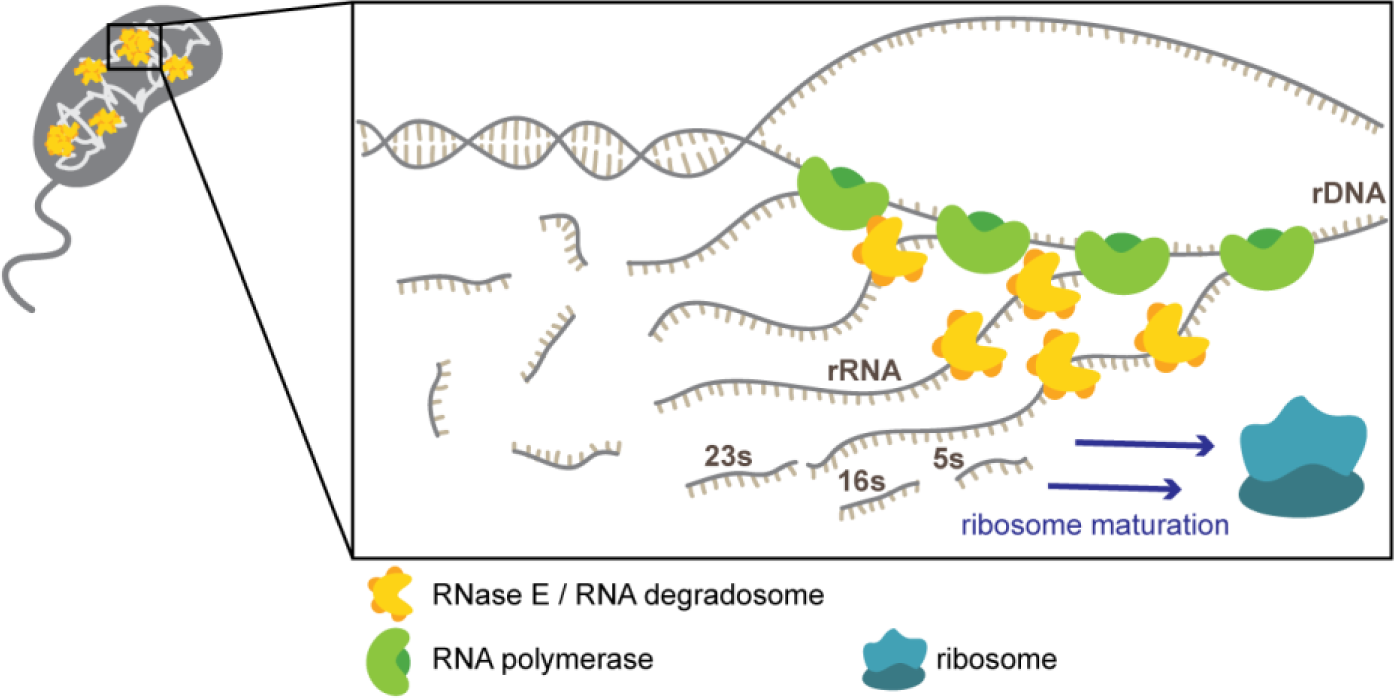
Model of RNase E’s integral role in RNA metabolism and ribosome biogenesis. RNase E clusters are found in the cytoplasm of the cell. A zoom in of a cluster co-localizing with an rDNA locus shows many RNAPs transcribing rRNA. These rRNAs are transcribed together in an operon and are cleaved by RNases, including RNase E, to generate mature rRNA species that are then assembled to form mature ribosomes.

## Methods

### Preparation of cell strains for imaging

*Caulobacter* cells were grown overnight from frozen culture in 5 mL of PYE growth medium^40^ with shaking in a 28°C water bath. The day before imaging, cultures were diluted 1:1000 into the defined minimal media M2G.^41^ After growing to mid-log phase, 1-mL aliquots of cells were washed once with M2G by centrifugation (3 min at 8,000 RPM, 4°C (Eppendorf 5430R)) and the pellet was resuspended into clean M2G. Synchronized cells were prepared using a Percoll density gradient,^42^ allowing the cells a 5 min recovery in M2G at 28°C before moving on to the next step. Rifampicin was added at 50 μg/mL final concentration (from 10 mg/mL stock solutions in DMSO (Fisher)) and incubated with the cells for 30 minutes with shaking. The cells were washed by centrifugation (3 min, 8000 RPM, 4°C (Eppendorf 5430R)) and resuspension in clean M2G. Cells were fixed with the addition of 1% formaldehyde (Fisher Scientific, Lot 091161), followed by a 10 min incubation at room temperature and 30 min incubation on ice, and finally washed three times with clean M2G.

Before imaging, the cells were resuspended in a small amount (~20-50 μL) of minimal medium, producing a concentrated cell suspension. To this suspension, ~1 nM of fiducial markers were added (Molecular Probes, 540/560 carboxylate-modified FluoSpheres, 100 nm diameter), then 1-2 μL of this mixture was deposited onto an agarose pad (composed of 1.5% (w/w) of low melting point agarose (Invitrogen) in M2G) and mounted onto an argon plasma-etched glass slide (Fisher) and imaged immediately.

For imaging the tetO/tetR labeled strains, the cells were incubated with 0.03% xylose final concentration (from 30% (w/v) stock solutions in water) for 30 minutes before washing once with M2G, then resuspending in M2G (~20-50 μL) and imaged immediately on an agarose pad as described above.

### Single-molecule imaging for super-resolution microscopy and single-particle tracking

Both SPT and SR microscopy rely on localizing single molecules with high precision; SPT is based upon localizing the same molecule over time to gain information about this molecule’s dynamics. In contrast, SR microscopy localizes many different SMs over time and then builds up a reconstruction to gain information about the static structure of interest. Since the imaging time required to acquire enough SM localizations for SR imaging can be long (~minutes), the structure of interest must be stationary on the time scale of imaging.

Cells immobilized on agarose pads were imaged on a home-built epifluorescence microscope (Nikon Diaphot 200) with additional optics inserted before the 512x512 pixel Si EMCCD camera (Andor Ixon DU-897), which enables two color, three-dimensional super-resolution imaging described previously.^43^ Fluorescence emission from the sample was collected through a high NA oil-immersion objective (Olympus UPlanSapo 100x/1.40 NA), filtered by a dual pass dichroic mirror (Chroma, zt440/514/561rpc), a 560 nm dichroic beamsplitter (Semrock FF560-FDi01), a 514 nm long pass filter (Semrock LP02-514RE), a 561 nm notch filter (Semrock NF03-561E) and a bandpass filter (Semrock, FF01-532/610). Two-dimensional white light transmission images of the cells were recorded before imaging. The eYFP was pumped with 514 nm excitation (Coherent Sapphire 514-100 CW, ~0.5-1 kW/cm^2^). A 405 nm laser (Obis, ~0.1-10 W/cm^2^) was used to photoactivate and a 561 nm laser (Coherent Sapphire 561-100 CW, ~0.5 kW/cm^2^) was used to excite fluorescence from PAmCherry. Laser intensity was determined by measuring the power at stage with Newport power meter (model 1918-C) and estimating the FWHM spot size by fitting the average fluorescence from the sample to a 2D Gaussian by nonlinear least squares using a custom MATLAB (Mathworks) program. Integration times were25 ms for SPT data and 50 ms for SR data.

### Data analysis procedure

#### Fitting 3D single molecule data

Both 3D SR and SPT data were fit using a modified version of Easy-DHPSF,^44^ which is freely available at https://sourceforge.net/projects/easy-dhpsf/. Calibration scans were created by axially scanning a nanohole array (NHA)^45^ filled with ~25 μL of ~50 μM Alexa 568 (Life Technologies) and Atto 520 (Sigma) dyes using a piezoelectric objective scanner (Mad City Labs, C-focus). These calibration scans were used to relate angular lobe orientation to axial position, and as templates to locate candidate single molecules in the data. The NHA allowed us to account for field-dependent errors arising from the differing rotation rates, and hence apparent axial position, of the DH-PSF as a function of lateral position. The background photons (~16-22 photons/pixel) were estimated using a median filter in Matlab.^44^ On average, we detected 1327 ± 449 photons per 25 ms frame for the eYFP-labeled RNase E and ribosomes. Localization precision was estimated with an empirical formula derived from repeatedly localizing single beads under variable background conditions.^43^ Images included in this paper have typical average localization precisions of 32 ± 8 nm (error is standard deviation), 33 ± 9 nm, and 49 ± 13 nm in *x, y,* and *z.* Systematic errors from sample drift resulting from mechanical and thermal fluctuations were corrected by adding ~1-10 nM concentration of bright 560/580 Fluospheres (Life Technologies) to the sample and using these as fiducial beads. High frequency noise in the drift is removed with wavelet filtering in Matlab.^43, 44^

#### Two-channel registration

Arrays of control point positions were generated by axially scanning the NHA^45^ filled with ~25 μL of ~50 μM Alexa 568 (Life Technologies) and Atto 520 (Sigma) dyes using a piezoelectric objective scanner (Mad City Labs, C-focus) and imaging the NHA simultaneously in the two channels using both the 514 nm and 561 nm lasers. The red channel was transformed to the green channel using the same algorithm described previously.^43^ TRE’s ranged from 5-15 nm and FRE’s ranged from 4-12 nm.

#### Removing oversampling

Oversampling, caused by including multiple localizations of the same protein from fluorescent protein labels that emit for more than one frame, creates spurious clustering. Before analyzing clustering in fixed cells, oversampling was removed with a radial and temporal filter. Localizations within a set radial and time region are assumed to belong to the same molecule and are combined into a single localization located at the average position. A reasonable radial distance was determined by simulating SR data containing oversampling on the size scale of the localization precision. The threshold radius (as a function of the mean localization precision) was varied until a grouping algorithm in Matlab recovered the ground truth number of simulated single molecules (thus, estimating the grouping of the oversampled clusters). The optimized spatial threshold was found to be 1.5*mean localization precision 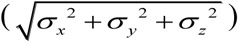. The temporal threshold was determined individually for each cell dataset by calculating the probability of finding another localization within the spatial threshold as a function of time. The probability typically followed a single exponential decay. The time constant of the exponential was used as the temporal threshold.

#### Diffusion analysis

Trajectories were constructed by connecting localizations from consecutive frames. Only trajectories of at least 4 steps (5 frames or 125 ms) were used for MSD analyses. The maximum allowed *xyz* displacements of RNase E-eYFP and L1-eYFP molecules over 25 ms were set to 1000 nm. Diffusion coefficients were calculated from ensemble-averaged 3D MSDs of individual RNase E molecules using the first two time lags to avoid long-time effects of confinement and non-Brownian motion. The reported errors are the standard errors of the mean diffusion coefficients as determined by bootstrapping (50% of the trajectories were sampled 500 times). Error bars in the MSD curves are SEM’s.

#### Simulating trajectories inside cell-shaped object

Brownian simulations were performed inside a cylindrical cell of radius 250 nm and length 3.5 μm with two hemispherical caps; finite sampling from a camera and motion blur were taken into account by averaging 10 positions for every 25 ms frame. Each 3D position was then given a *xyz* “kick” from a Gaussian distribution to simulate localization precision.

#### Confinement analysis

To test for confinement, we divided each trajectory (of at least 4 steps (5 frames or 125 ms) in length) into sub-trajectories of 4 steps (5 frames or 125 ms). For each sub-trajectory, the maximum displacement from the first point of that sub-trajectory was calculated.^20, 21^ We performed simulations of Brownian diffusers inside a cylindrical cell volume with hemispherical caps to determine whether a sub-trajectory was confined or not. The effect of confinement, such as in the case of cytoplasmic molecules diffusing in a cell, results in a smaller value for the apparent, observed diffusion coefficient than in the case in which the molecule is not confined by cell walls. To ensure that we were making a valid comparison of diffusion coefficients in our confinement analyses,^46^ we performed Brownian simulations of a cytoplasmic molecule inside a cylindrical cell with hemispherical caps. We used an empirical diffusion coefficient that resulted in an apparent diffusion coefficient approximately equivalent to the ensemble-averaged measured diffusion coefficients for WT and rif-treated cells (Supplementary Fig. S4). A threshold was set such that only 5% of the simulated Brownian sub-trajectories are confined. Sub-trajectories with displacements less than this value (118 nm for WT and 123 nm for rif-treated, plotted as vertical dotted lines in Fig. 1e) were classified as confined. From the cumulative distributions shown in Fig. 1e, this criterion would imply that ~50% of the WT sub-trajectories were mobile. However, we found that many molecules in this group did not diffuse very far, as suggested by fixed snapshots of the spatial distribution (Fig. 3e). To include the possibility of an intermediate degree of confinement, another threshold was arbitrarily set at the Brownian 50^th^ percentile, and subtrajectories with displacements greater than this value (198 nm for WT and 208 nm for rifampicin-treated) were classified as clearly mobile. All other sub-trajectories were classified as intermediate.

#### Fitting squared displacement CDF’s to multiple populations

To extract the different populations of ribosomal L1 proteins, we fit the empirical CDF of the squared displacements of our trajectories to Eqn. (1),^47^ where *r*^2^ are the experimental squared displacements, *τ* is the time lag, *x_i_* is the weight of population *i*. Eqn. (2) (in which *d* is the number of dimensions, *D_i_* is the diffusion coefficient of population *i*, Δ*t* is the camera exposure time, σ_*j*_ is the localization sigma in the j^th^ dimension) is substituted into Eqn. (1) to perform the fit. The fitting is constrained by Eqn. (3) to ensure that all weights sum to 1. The fitting is performed in Matlab, minimizing the objective function (*P_data_ – P*_*fit*_)^2^ using the function fmincon.

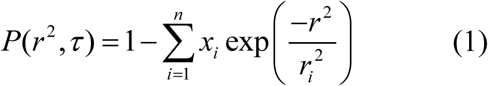

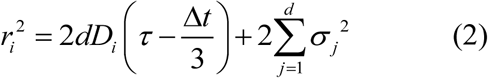

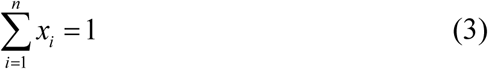

Fitting with three populations yields the same result as fitting with only two populations (Supplementary Fig. S9).

#### Clustering analysis by spatial point statistics

To quantify the degree of clustering in our SR images, we used spatial point statistics. Complete spatial randomness (CSR) was simulated within each cell volume using an identical number of points with a custom MATLAB (Mathworks) program. The simulated points were then thinned to be the same number of localizations as the data, then a random “kick” of localization precision error (pulled from a Gaussian distribution of the mean localization precision of the data) in all three dimensions to each simulated point. In all cases, 100 simulations were performed, as this balanced computational time with statistical information.

The uncertainty in the data from the localization precision was also bounded by simulation. Each detected localization had its own localization precision error. A cloud of points (100 simulations) with shape dictated by the individual localization precision were simulated around each localization. The same statistical metrics were calculated on these simulated points, and the 95% confidence bounds were used to report the error in the data from the localization precision.

Global clustering is quantified using Ripley’s K-function^29^ and its normalized forms.^48^ As previously described, Ripley’s K-function represents the average probability of finding localizations a distance r from the typical localization.^49^ For CSR, K(r) scales with the volume and is described by Eqn. 4 where *b_d_r^d^* is the volume of the unit sphere in d dimensions. Deviations from this shape indicate clustering or repulsion; however, visualizing deviations from a straight or flat line is simpler than for an exponential curve. The normalized K-function is given by L(r) (Eqn. 5), which can further be normalized into the H(r) function, such that CSR is a flat horizontal line at *H =* 0 (Eqn. 7).^50^ To be able to analyze clustering across many cells, the quantity shown in Eqn. 7 was calculated for each cell.

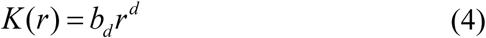

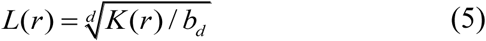

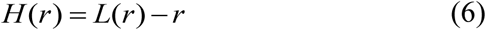

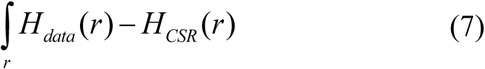

#### Clustering analysis by density threshold

For the 3D two-color RNase E/PopZ datasets, we use a density-based clustering algorithm, density-based spatial clustering of applications with noise (DBSCAN),^51^ which detects clusters according to the local density of localizations within a certain search radius. For the RNase E clusters, we used the parameters ε = 70 nm and minPts = 6; for the PopZ clusters, we used the parameters ε = 200 nm and minPts = 7. These thresholds were chosen empirically.

In the RNase E-eYFP/PAmCherry-PopZ cells, the clustering results for RNase E did not give an absolute count of the number of molecules in a cluster, but rather gave the relative amounts of number of molecules detected. This is due to the photophysical properties of eYFP. Imaging SMs of eYFP requires an initial bleachdown of the sample, then waiting for the stochastic recovery and blinking of this FP. The ideal situation is one in which each copy of eYFP recovers. However, in practice, only about 5-15% of the molecules recover and are detected.

#### Super-localization of labeled loci and colocalization

50 frames (100 ms exposure) were summed for tetR-eYFP and RNase E-PAmCherry. TetR-eYFP loci were super-localized by fitting 2D Gaussians (which well-approximates the point spread function of the microscope) in ThunderSTORM (ImageJ). These localizations were then transformed on to the RNase E-PAmCherry images using the mapping function generated with the following method. The array of 500 control points that were detectable in both channels simultaneously and evenly distributed across the whole field of view were localized by fitting a 2D Gaussian in ThunderSTORM. These localizations were used to generate a mapping function between the two channels using local weighted mean transformation in custom MATLAB scripts to take into account field dependence of the transformation. Percentage of DNA loci co-localizing with an RNase E cluster was determined by visual inspection of the transformed locus with respect to the RNase E diffraction-limited image.

## Acknowledgements

The authors thank the James Weisshaar lab at University of Wisconsin-Madison for sending the *E. coli* S2 ribosome strain, Melanie Barnett and Jillynne Quinn in the lab of Sharon Long, Dept of Biology, Stanford University for help creating the ribosome *S. meliloti* strain, Thomas H. Mann for help with constructing the RNase E-eYFP/PAmCherry-PopZ strain, and Alex von Diezmann for the Brownian diffusion simulation code. This work was supported in part by the National Institute of General Medical Sciences, Grants No. R01-GM086196 to W.E.M. and L.S., R35-GM118067 to W.E.M., R35-GM118071 to L.S. L.S. is a Chan Zuckerberg Biohub Investigator and J.M.S. was an NIH Postdoctoral Fellow No. F32-GM100732.

## Author contributions

C.A.B., J.W., M.K.L., J.M.S., L.S. and W.E.M. conceived and designed the experiments. C.A.B., J.W., and J.M.S. made bacterial strains. C.A.B., J.W., and M.K.L. performed experiments. C.A.B., J.W., M.K.L., and W.E.M. analyzed the data. C.A.B., J.W., L.S., and W.E.M. wrote the manuscript.

The authors declare no competing financial interests.

## Supplementary methods

### Construction of fluorescently-labeled cell strains

#### JS51 NA1000 RNase E-YFP

RNase E-YFP fusion was generated by cloning the last 534bp of the RNase E gene (CCNA01954) lacking a stop codon into pEYFPC-1.^52^ First the last 500bp of the RNase E gene were amplified by PCR from the NA1000 genome using the following DNA primers:

CCNA01954 F aaacatatgccgagcgggccgagcgcg
CCNA01954 R tatggtaccccggcgccaccagccccgacg

Next, the resulting PCR product was digested with NdeI and KpnI enzymes, column purified, then ligated to NdeI/KpnI cut pEYFPC-1 plasmid. The ligation was transformed into *E. coli* top10 cells and selected on LB-spec plates. Resulting specR colonies were then screened by PCR for the insert, and then verified by Sanger sequencing (GENEWIZ). The purified plasmid was then transformed into NA1000 cells via electroporation and plated on PYE-spec plates. Resulting colonies were verified to be expressing the YFP fusion by fluorescence microscopy and the protein product was verified by western blot.

#### JS289 NA1000 RNase E-PAmCherry

RNase E-PAmCherry fusion was generated by cloning the last 534bp of the RNase E gene (CCNA01954) lacking a stop codon into pPAMC-4. First the last 500bp of the RNase E gene were amplified by PCR from the NA1000 genome using the following DNA primers:

CCNA01954 F aaacatatgccgagcgggccgagcgcg
CCNA01954 R tatggtaccccggcgccaccagccccgacg

Next, the resulting PCR product was digested with NdeI and KpnI enzymes, column purified, then ligated to NdeI/KpnI cut pPAMC-4 plasmid. The ligation was transformed into *E. coli* top10 cells and selected on LB-gent plates. Resulting gentR colonies were then screened by PCR for the insert, and then verified by Sanger sequencing (GENEWIZ). The purified plasmid was then transformed into NA1000 cells via electroporation and plated on PYE-spec plates. Resulting colonies were verified to be expressing the PAmCherry fusion by fluorescence microscopy.

#### JS290 NA1000 L1-eYFP

The ribosomal protein L1-eYFP fusion was generated by cloning the 3′ 500bp of the gene lacking a stop codon into pEYFPC-4 by Gibson assembly^52^ resulting in pL1::YFPC-4. First, the L1 fragment was generated by PCR from NA1000 cells using the L1-frag primers, and the pEYFP plasmid was linearized by PCR using the pEYFPC-4 primers. Next, PCR fragments were gel purified and equimolar concentrations of each were mixed in Gibson assembly mixture. The resulting pL1::YFPC-4 plasmid was screened for gentamicin resistance and sequence verified. The resulting pL1-YFPC-4 plasmid was then transformed into NA1000 cells by electroporation, and the resulting colonies were screened using fluorescence microscopy and the protein product was verified by western blot.

(L1-frag)_forward AATTAATATGCATGTCGAAGCCGCCATCGCC
(L1-frag)_reverse ATCTTAAGGTACCGGCGCCGATCGAGCTGATG
(pEYFPC-4)_forward CTCGATCGGCGCCGGTACCTTAAGATCTCGAGCTCC
(pEYFPC-4)_reverse GATGGCGGCTTCGACATGCATATTAATTAAGGCGCCTGC

#### RNase E-eYFP/PAmCherry-PopZ

The two color imaging strain of RNase E/PopZ was constructed by transducing JS51 with lysate from JP463 [popZ⸬PAmCherry-PopZ].

#### RNase E-eYFP in AHU2 background

An HU2 deletion strain MS087 (unpublished) that harbors HU2⸬aadA was transfected with lysate from strain JS51.

#### JS54 S. meliloti L1-eYFP strain

*S. meliloti* ribosomal protein L1-YFP plasmid psmeL1⸬YFPC-1 was generated by PCR amplifying the 3’ region of the gene lacking a stop codon with the following primers:

L1 Smeliloti F (PacI) ATT AATT AACGAACGGCACCGGCCG
L1 Smeliloti R (KpnI) AAGGTACCGGCAGCGCTGATCGTTGCCG

Resulting PCR product was then gel purified, cut with PacI and KpnI, then ligated into pEYFPC-1 cut at the PacI and KpnI sites. The resulting plasmid containing an in-frame eYFP was then sequence verified and mated into Sme 1021 and selected on M9-sucrose plates supplemented with 200 μg/mL spectinomycin. Resulting colonies were re-struck two additional times on M9-sucrose-spec plates.

#### tetO/tetR-labeled strains

TetR-eYFP was expressed under xylose promoter in *Caulobacter* from an integrative plasmid (LS5037 or pHPV472).^33^ To construct plasmids for labeling the two rRNA loci (CCNAR0069-CCNAR0064, CCNAR0087-CCNAR0082) with (TetO)s5 but avoid perturbing terminator sequences, 800 bp downstream genes CCNA02705-CCNA02706 and CCNA03637, respectively, were amplified by PCR; (TetO)s5 was amplified from pBS-85xtetO-KlURA3,^53^ both of which were then cloned into pMCS-2^52^ multiple cloning site yielding plasmids pJW349 and pJW350, respectively. To construct plasmids for labeling the DNA locus of CCNA01879 as our control, the last 800 bp of CCNA01879 was amplified and cloned into inverse PCR amplified plasmid backbone of pMCS-2 multiple cloning site yielding pJW404. Gibson assembly was used for all cloning.^54^ All resulting plasmids were then transformed into CB15N by electroporation.^41^

Clones in which the plasmids had integrated by single homologous recombination onto the chromosome were verified by PCR and were transduced with ΦCr30-dervived lysate^41^ from strain LS5037, a CB 15N derivative carrying the xyl ⸬TetR-eYFP, and then subsequently transduced with lysate from strain JS289 yielding strains JW523 [CCNA02705⸬(tetO)s5; xyl⸬TetR-eYFP; *RNase* E⸬RNase E-PAmCherry], JW524 [CCNA03637⸬(tetO)s5; xyl⸬TetR-eYFP; *RNase* E⸬RNase E-PAmCherry], JW525 [CCNA01879⸬(tetO)s5; xyl⸬TetR-eYFP; *RNase* E⸬RNase E-PAmCherry].

### Preparation of cell strains for imaging

#### Labeling cell surface amines with rhodamine spirolactam dye

Log-phase cells expressing RNase E-eYFP were washed by centrifugation and resuspension in clean M2G. Cells were fixed with the addition of 1% methanol (Fisher), followed by a 10 min incubation at room temperature then a 30 min incubation on ice, then washed three times with clean M2. The fixed cells were then resuspended in a mixture of 1 mL M2 and 50 μL of rhodamine spirolactam dye 9 (dissolved in DMSO (Fisher) at ~100 nM - 1 μM),^55^ then covered in foil and incubated at room temperature while shaking for 30 minutes. The cells were washed 5 times with clean M2, then resuspended in a small amount of M2 to produce a concentrated cell suspension. ~1 nM of fiducial markers were added (Molecular Probes, 540/560 carboxylate-modified FluoSpheres, 100 nm diameter), then 1-2 μL of this mixture was deposited on an agarose pad and imaged immediately.

#### Diffraction-limited imaging of labeled loci

Fluorescently-labeled DNA loci and RNase E were visualized by first imaging the tetR-labeled DNA loci with 514 nm laser (Coherent Sapphire 514-100 CW, ~0.05 kW/cm^2^) for 150 frames (100 ms exposure). RNase E-PAmCherry was then visualized by first activating PAmCherry with a 405 nm laser for 30 frames and then imaged with a 561 nm laser (Coherent Sapphire 561-100 CW, ~0.05 kW/cm^2^) for 150 frames.

---

**Figure S1:**
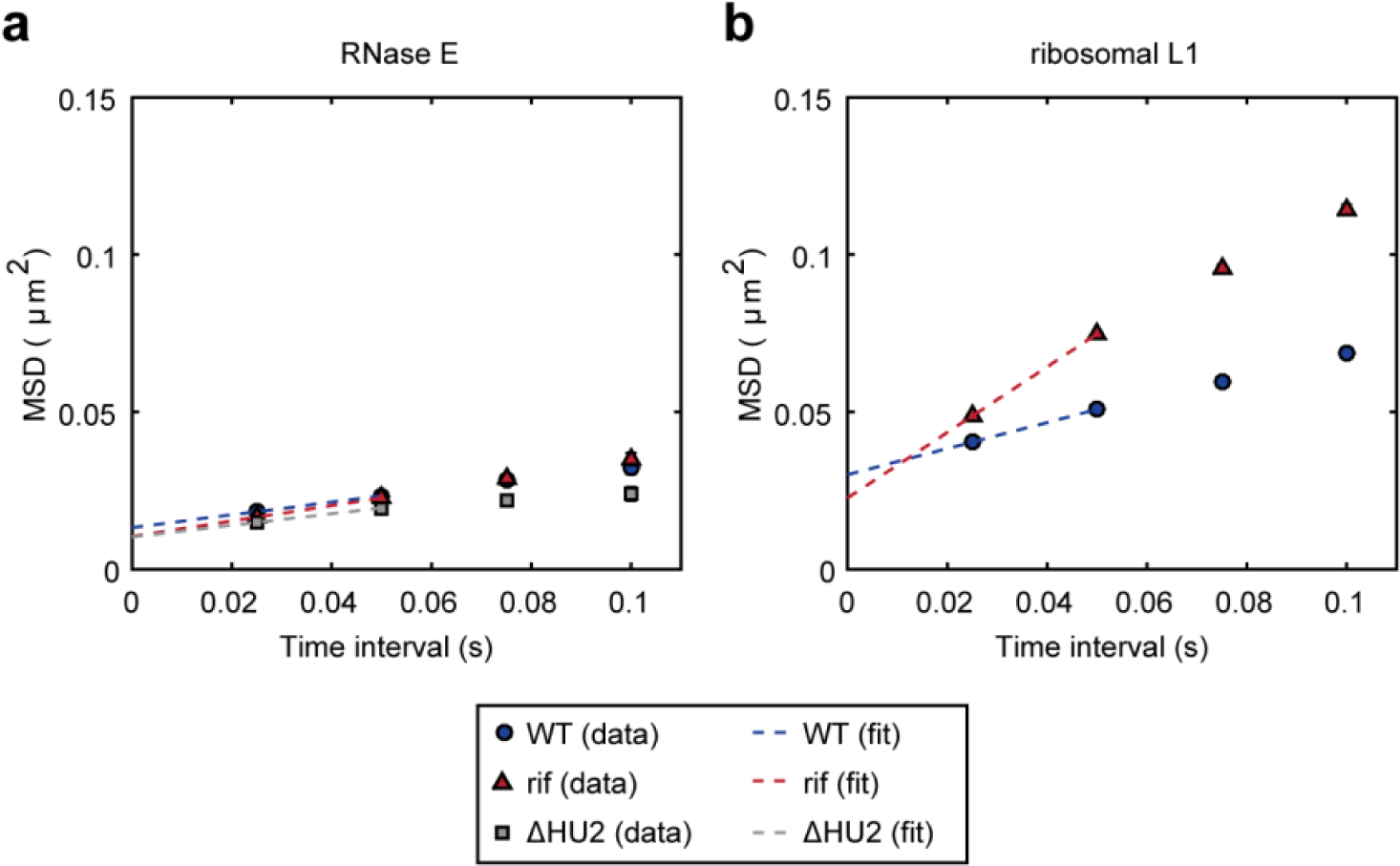
Ensemble-averaged MSD curves. **a,** Ensemble-averaged MSD curves for RNase E. **b**, Ensemble-averaged MSD curves for ribosomal-protein L1. Error bars are SEM’s.

**Figure S2:**
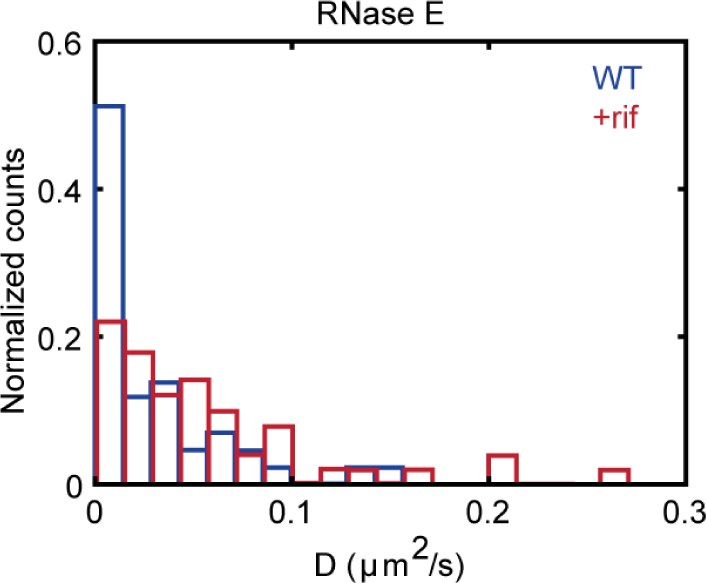
Distribution of RNase E SM diffusion coefficients for different trajectories.

**Figure S3:**
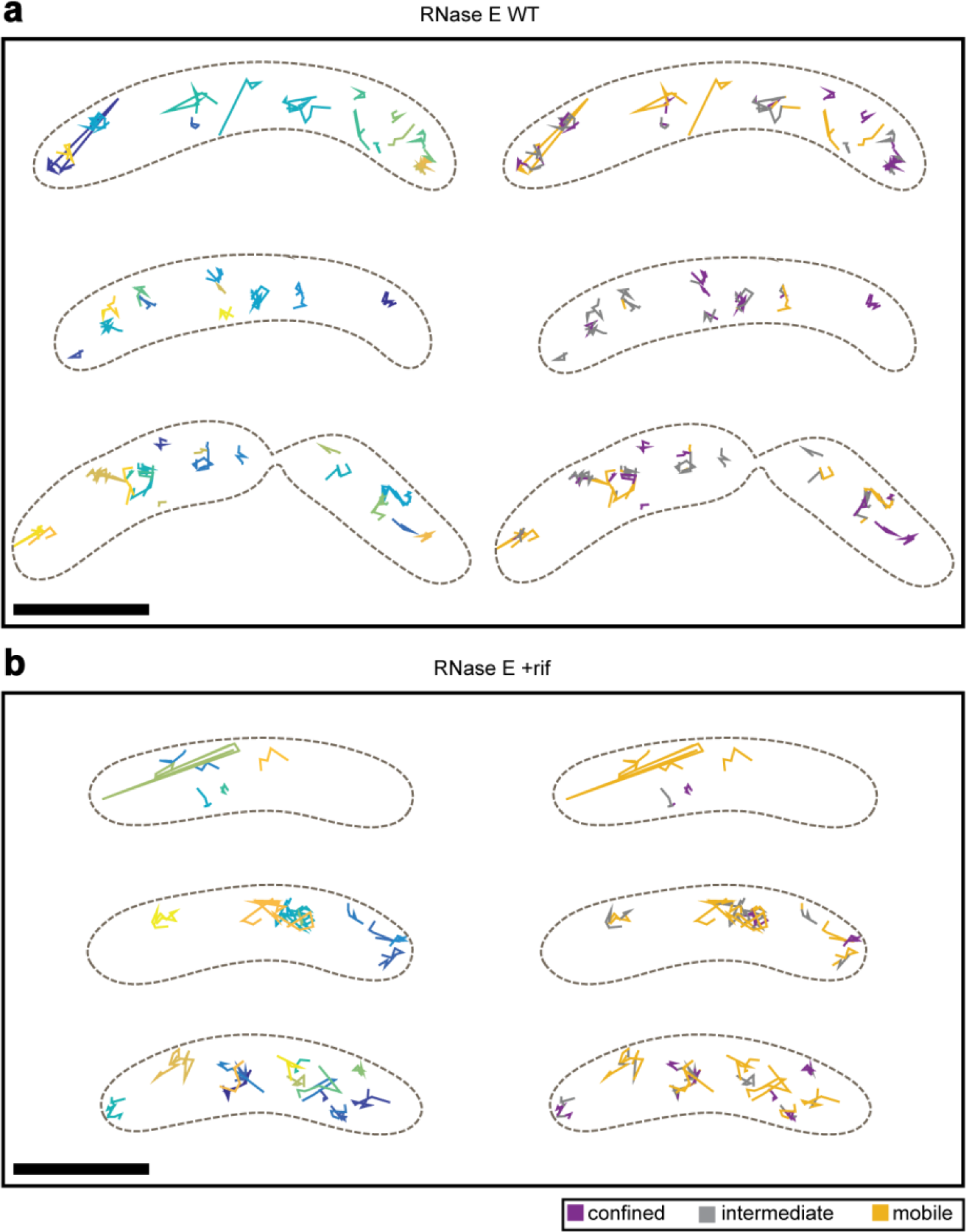
Additional examples of live *Caulobacter* cells and SPT trajectories of RNase E. **a**, Left, WT cells and trajectories, where each trajectory is plotted as a different color. **a,** Right, WT cells and each trajectory plotted according to its confinement level. **b,** Left, rif-treated cells and trajectories, where each trajectory is plotted as a different color. **b**, Right, rif-treated cells and each trajectory plotted according to its confinement level. Scale bars are 1 micron.

**Figure S4:**
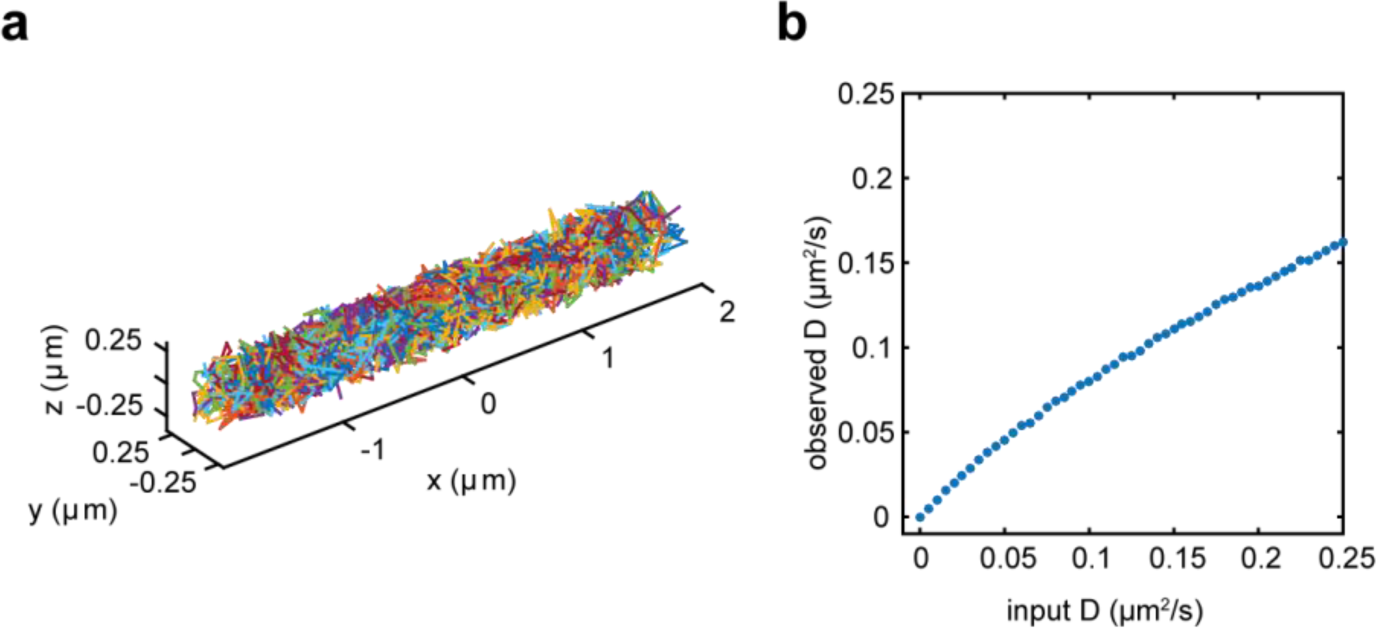
Results of Brownian simulations in a cylindrical cell volume with hemispherical caps. **a**, An example cell with a 250 nm radius and a 3.5 μm length with 500 trajectories. The input diffusion coefficient for this cell is 0.100 μm^2^/s. The simulation was performed with 9 steps (10 frames or 250 ms). Each trajectory is plotted as a different color. **b**, Observed D vs input D for simulations performed in the same volume as a. 5000 trajectories were simulated for each input D.

**Figure S5:**
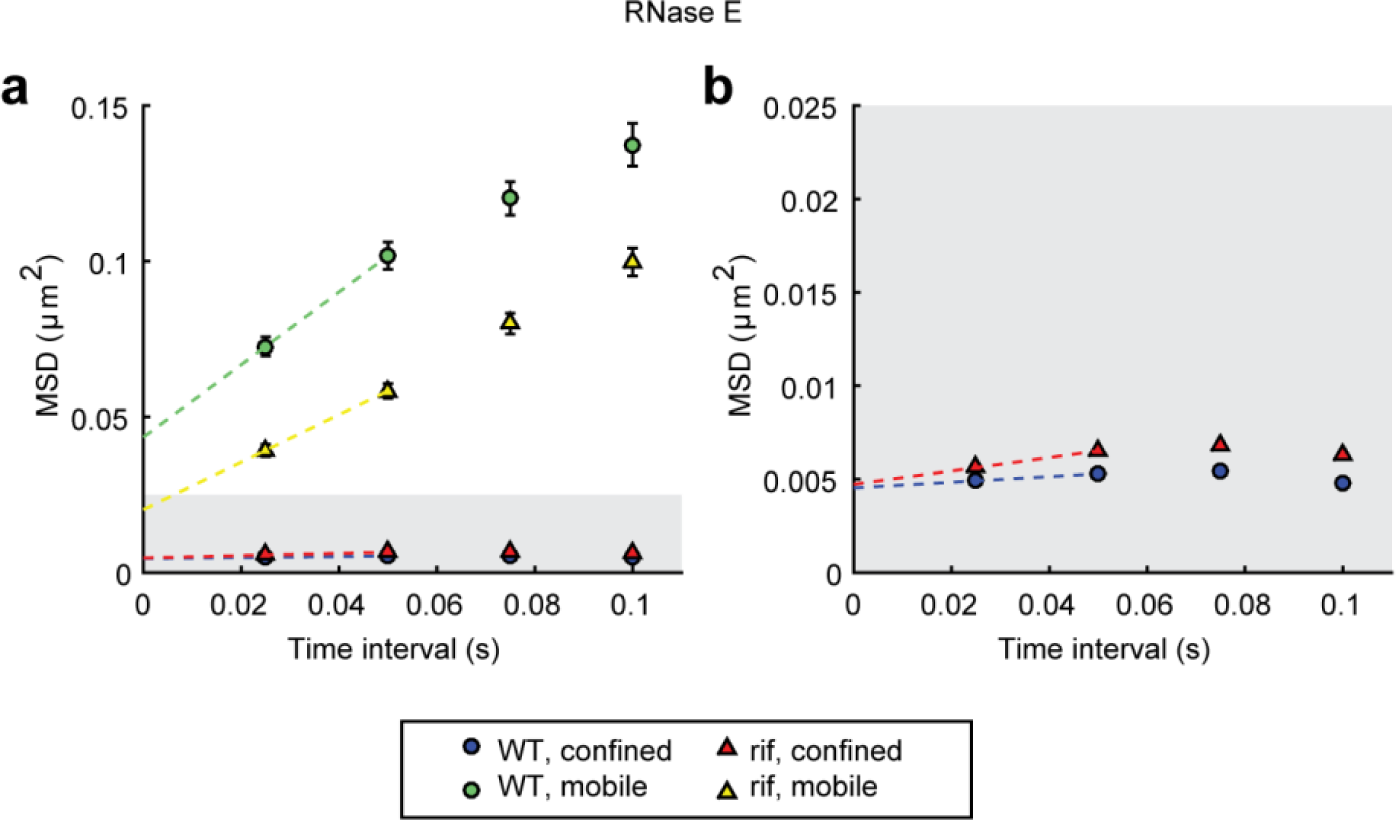
Ensemble-averaged MSD curves for confined and mobile RNase E subtrajectories. **a**, Ensemble-averaged curves for WT confined (blue circle) vs mobile (green circle) molecules and rif-treated confined (red triangle) vs mobile (yellow triangle) molecules. **b**, Zoom in of the gray region in a, showing greater motion of confined rif-treated subtrajectories than confined WT subtrajectories. Error bars are SEM’s.

**Figure S6:**
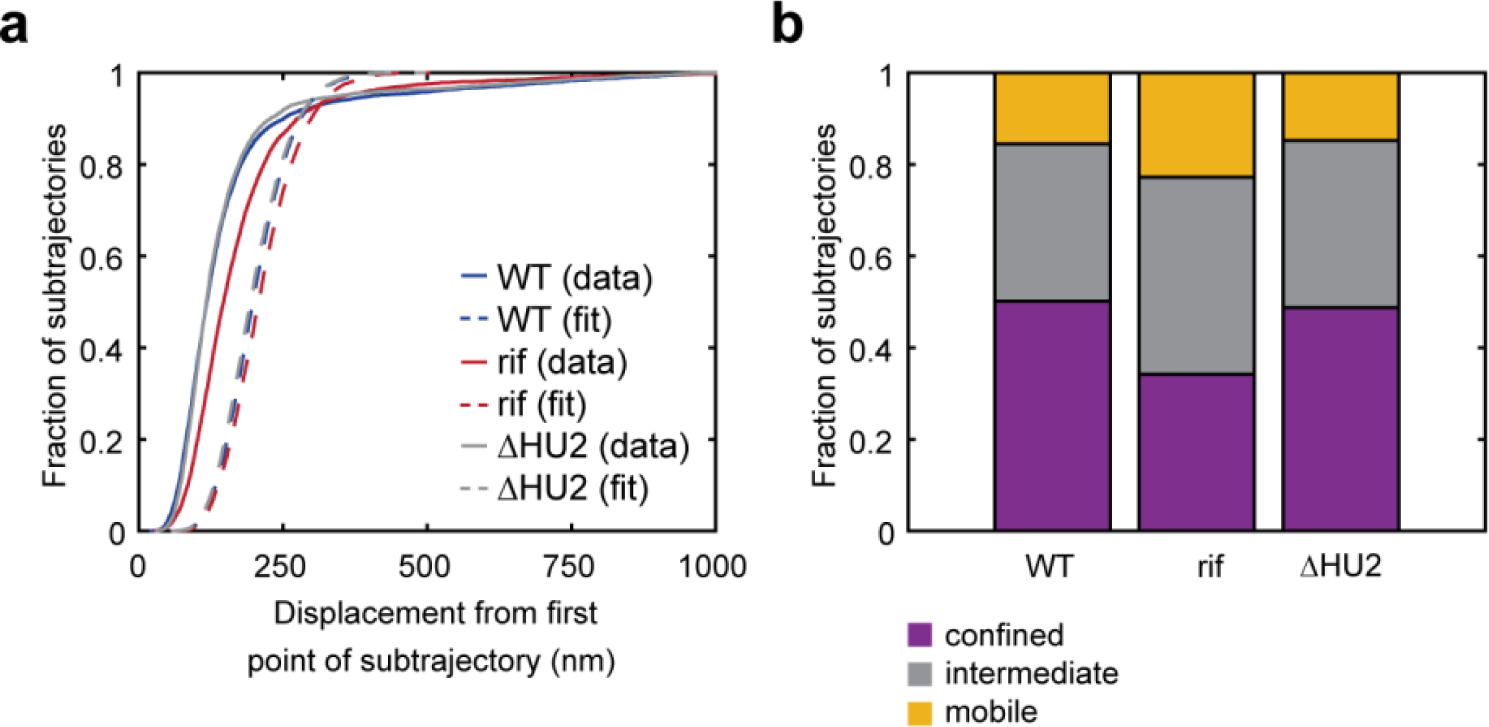
Results from the confinement analysis done for RNase E. **a**, CDFs from our confinement analyses. b, Fractions of confined, intermediate, and mobile for WT, rif-treated, and ΔHU2 cells. The fraction of confined subtrajectories for WT/rif/ΔHU2 cells are 50.1%/34.2%/48.7%. The fraction of mobile subtrajectories for WT/rifΔHU2 cells are 15.6%/22.8%/14.8%. Analysis was performed on 2107 trajectories from 270 HU2 deletion cells

**Figure S7:**
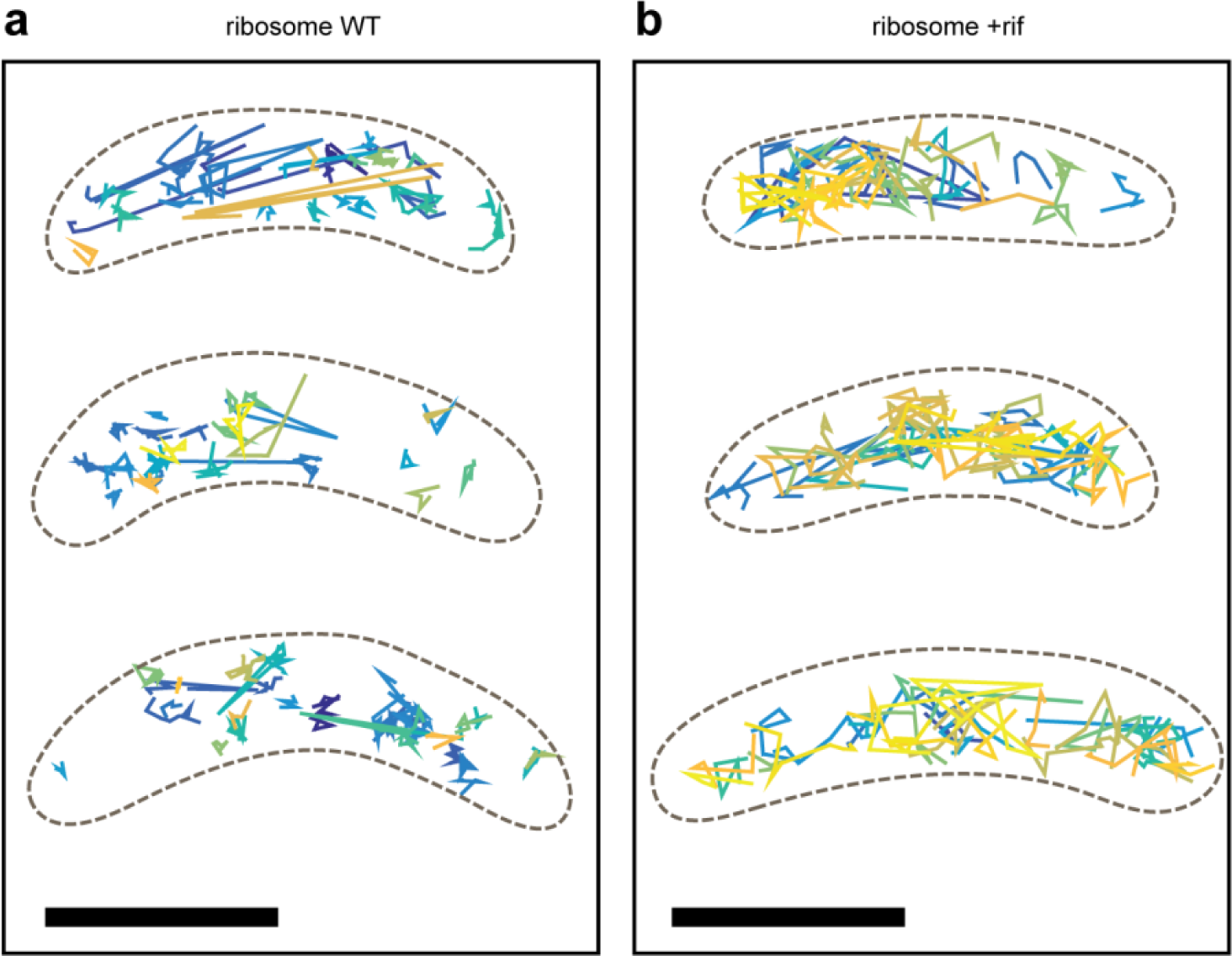
More example live *Caulobacter* cells and trajectories from SPT of ribosomal-associated protein L1. **a**, WT cells and trajectories, where each trajectory is plotted as a different color. **b**, Rif-treated cells and trajectories, where each trajectory is plotted as a different color. Scale bars are 1 micron.

**Figure S8:**
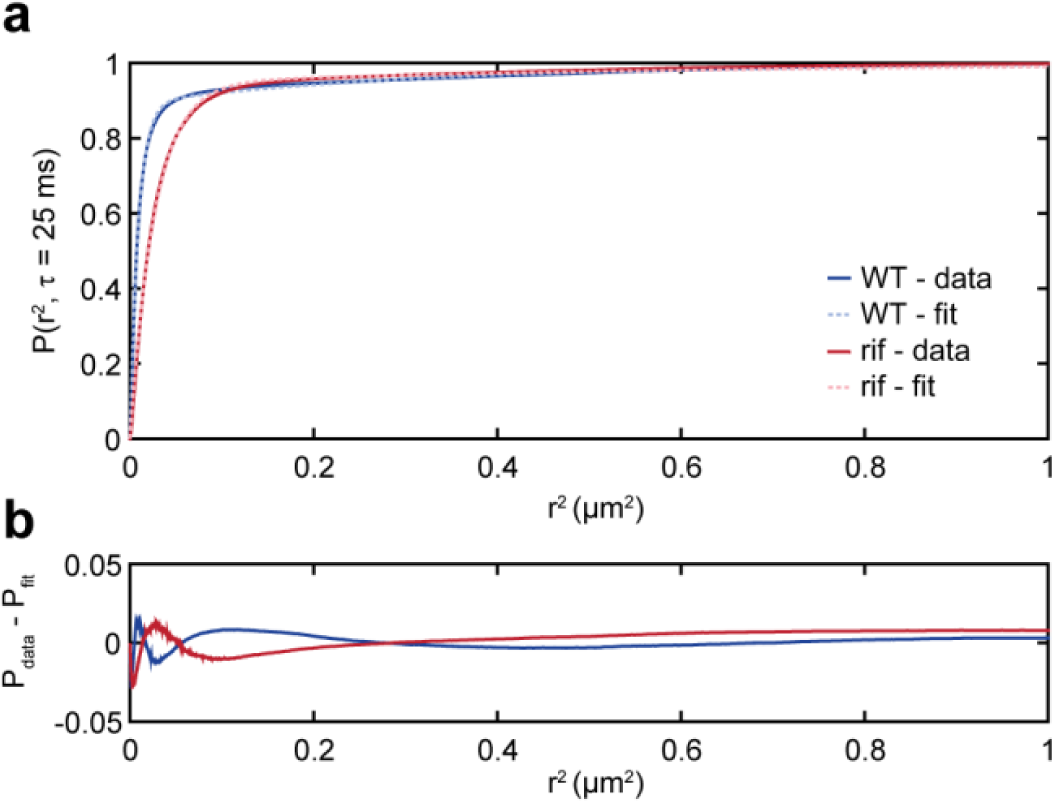
CDF of squared displacements of L1 trajectories fitting with two populations. **a**, CDF’s of squared displacements of L1 trajectories (solid lines) and their fit to a two-population model (dotted lines). **b**, Residuals from the fits.

**Figure S9:**
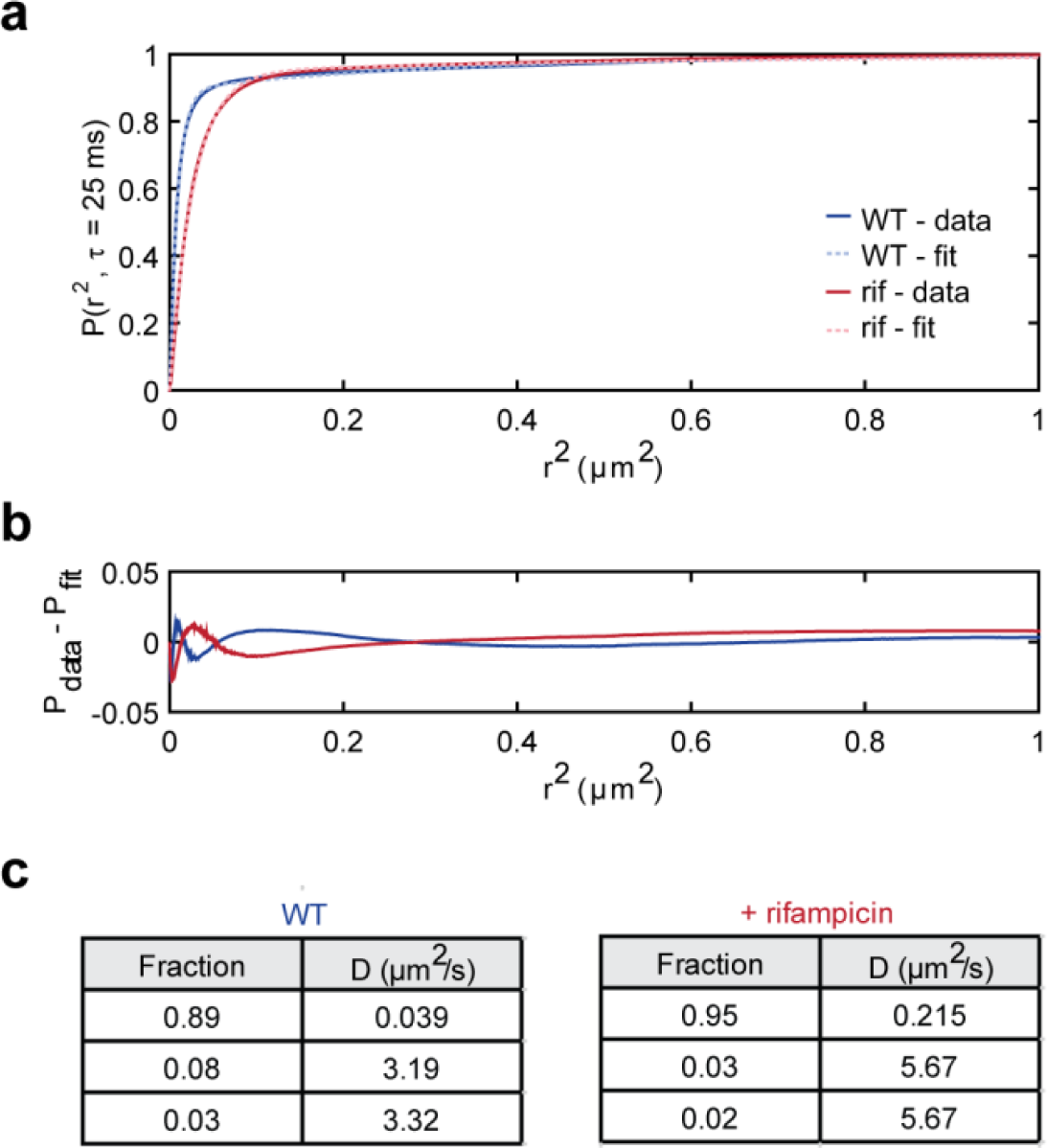
CDF of squared displacements of L1 trajectories fitting with three populations. **a**, CDF’s of squared displacements of L1 trajectories (solid lines) and their fit to a three-population model (dotted lines). **b**, Residuals from the fits. **c**, Summary of results from fitting to a three-population model.

**Figure S10:**
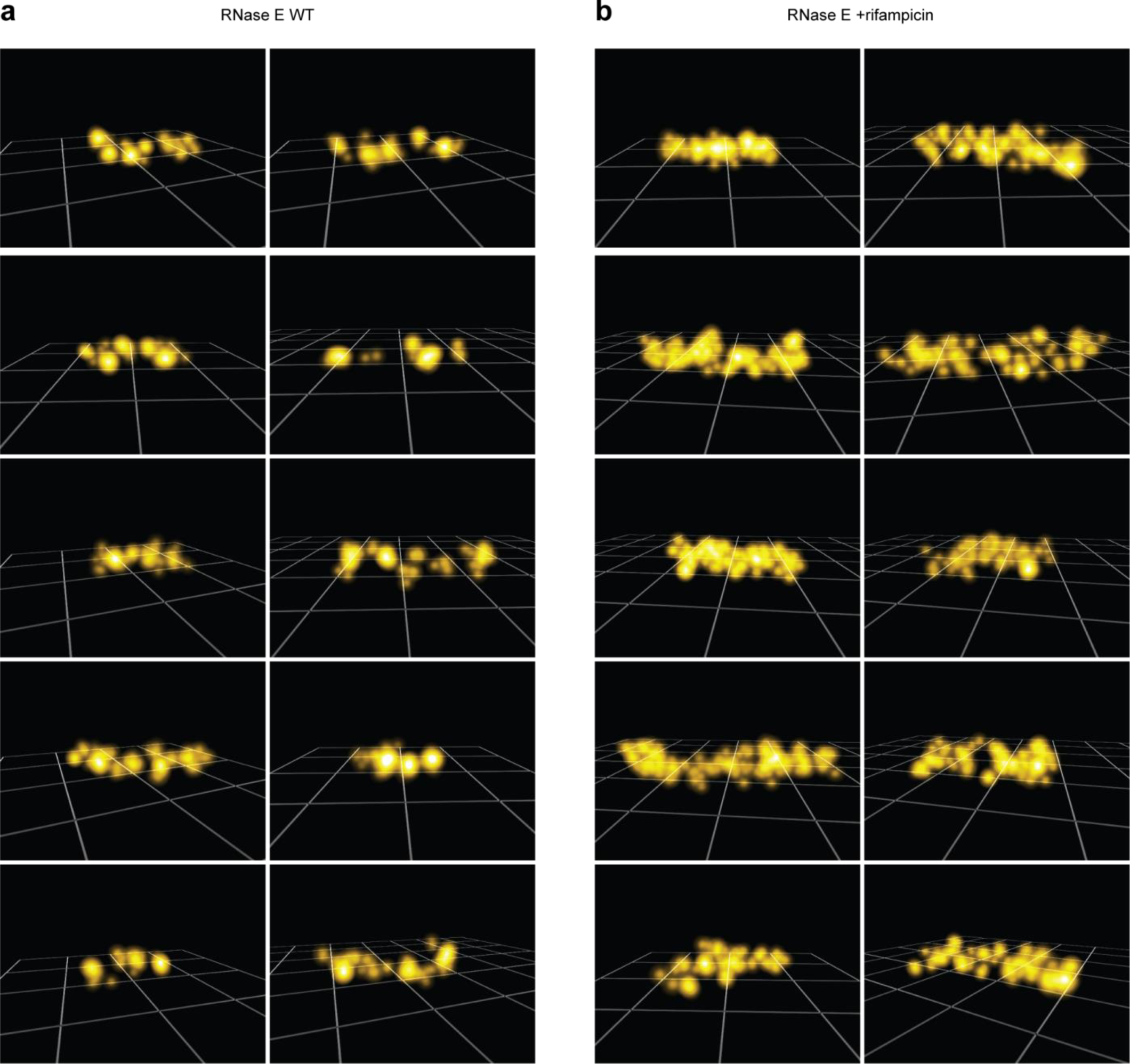
More examples of RNase E SR reconstructions. **a**, Examples of WT fixed cells expressing RNase E-eYFP. **b**, Examples of rif-treated fixed cells expressing RNase E-eYFP. Cells are on 1 micron grids. Each localization is plotted as a 3D Gaussian with a sigma equivalent to the average xy localization precision of the localizations in each cell (mean of 29/30/43 nm in *x/y/z* for all cells). On average, we detect 120 molecules per cell and 1557 photons per localization.

**Figure S11:**
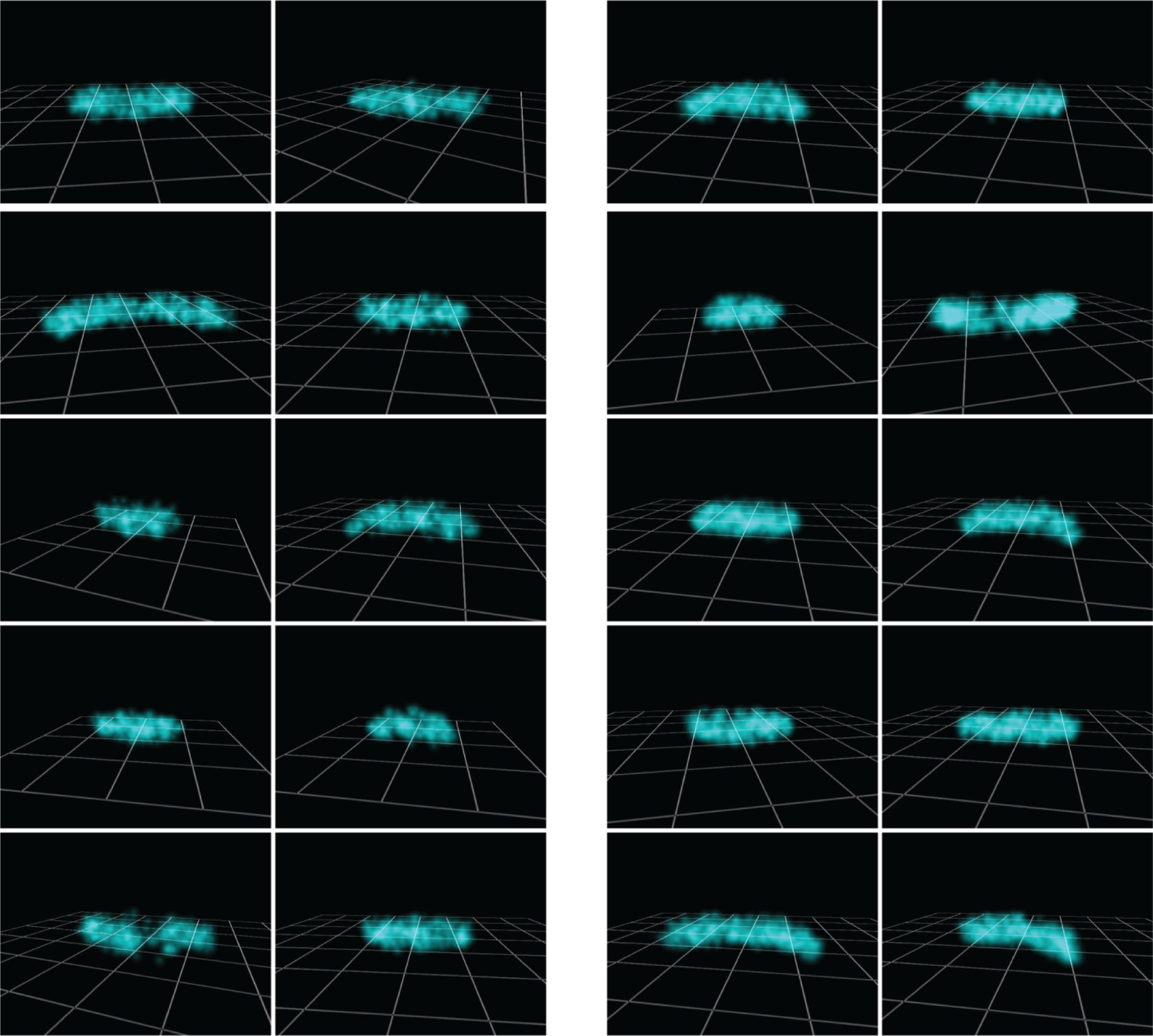
More examples of ribosomal-associated protein L1 SR reconstructions. **a**, Examples of WT fixed cells expressing L1-eYFP. **b**, Examples of rif-treated fixed cells expressing L1-eYFP. Cells are on 1 micron grids. Each localization is plotted as a 3D Gaussian with a sigma equivalent to the average xy localization precision of the localizations in each cell (mean of 24/25/37 nm in *x/y/z* for all cells). On average, we detect 720 molecules per cell and 2182 photons per localization.

**Figure S12:**
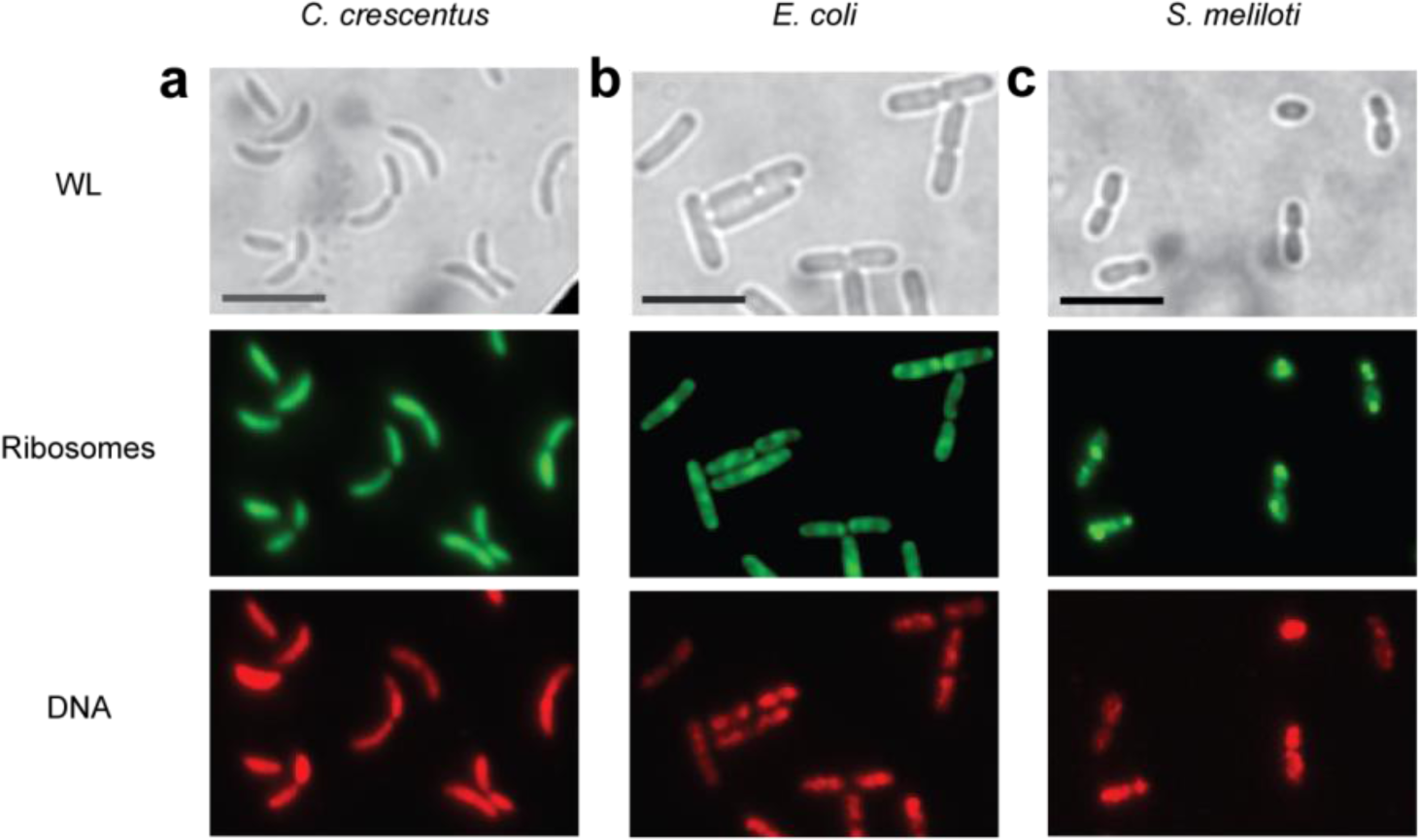
DL imaging of *C. crescentus, E. coli,* and S. *meliloti* ribosomes and DNA. a, top, WL image of live Caulobacter expressing ribosomal-associated protein L1-eYFP. **a**, middle, DL image of L1-eYFP in *Caulobacter*. **a**, bottom, DL image of *Caulobacter* DNA, visualized using the live-cell-permeable dye SYTOX orange. **b**, top, WL image of live *E. coli* expressing ribosomal-associated protein S2-eYFP. b, middle, DL image of S2-eYFP in *E. coli.* **b**, bottom, DL image of *E. coli* DNA, visualized using SYTOX orange. **c**, top, WL image of live *S. meliloti* expressing ribosomal-associated protein L1-eYFP. **c**, middle, DL image of L1-eYFP in S. *meliloti.* c, bottom, DL image of *S. meliloti* DNA, visualized using SYTOX orange.

**Figure S13:**
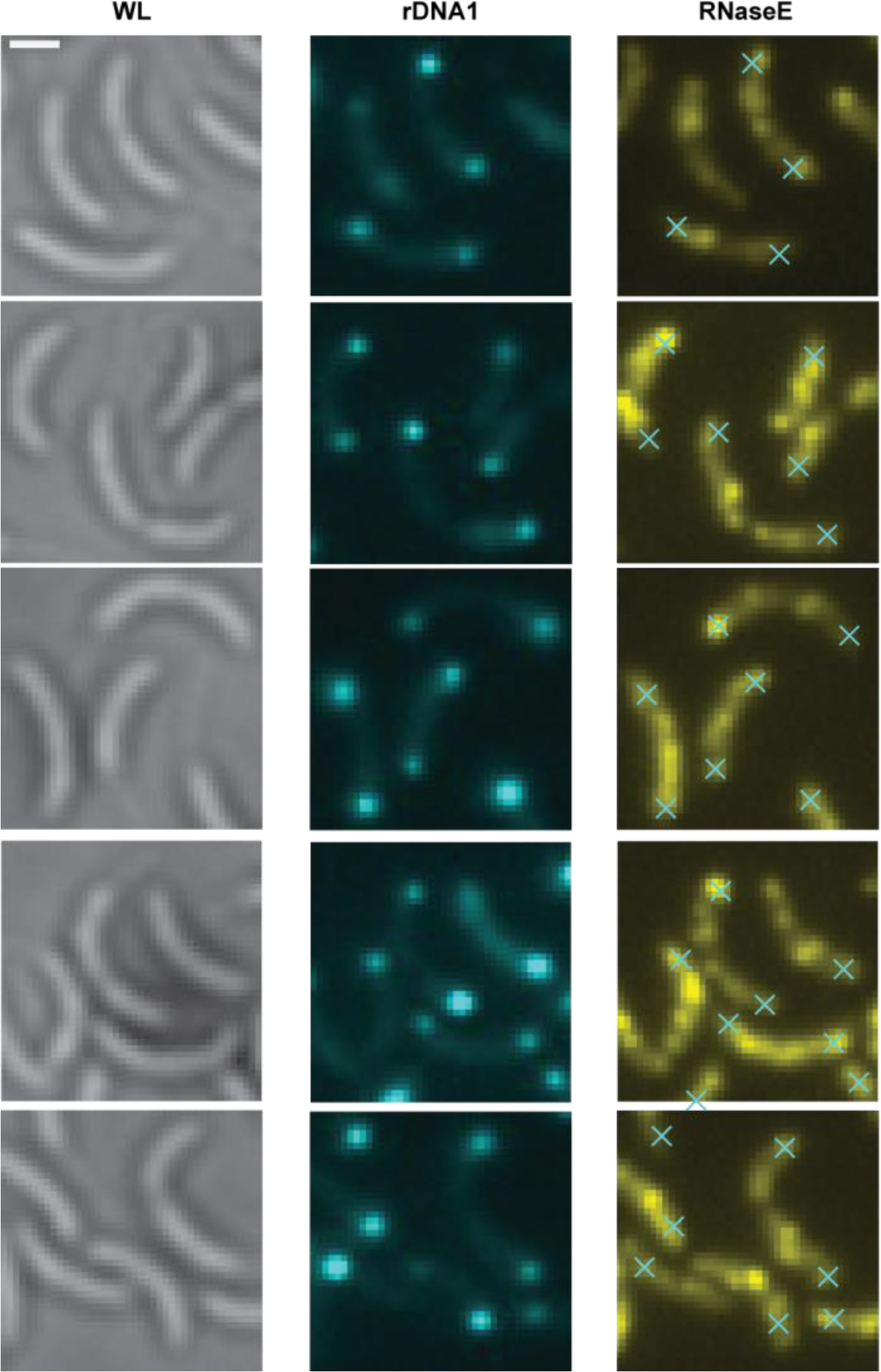
Additional DL examples of rDNA1 and RNase E colocalization in *Caulobacter*. Left column shows WL images of *Caulobacter* expressing RNase E (visualized in right column) and rDNA1-tetO/tetR (visualized in middle column). Crosses in right column show the locations of chromosomal loci displayed in middle column.

**Figure S14:**
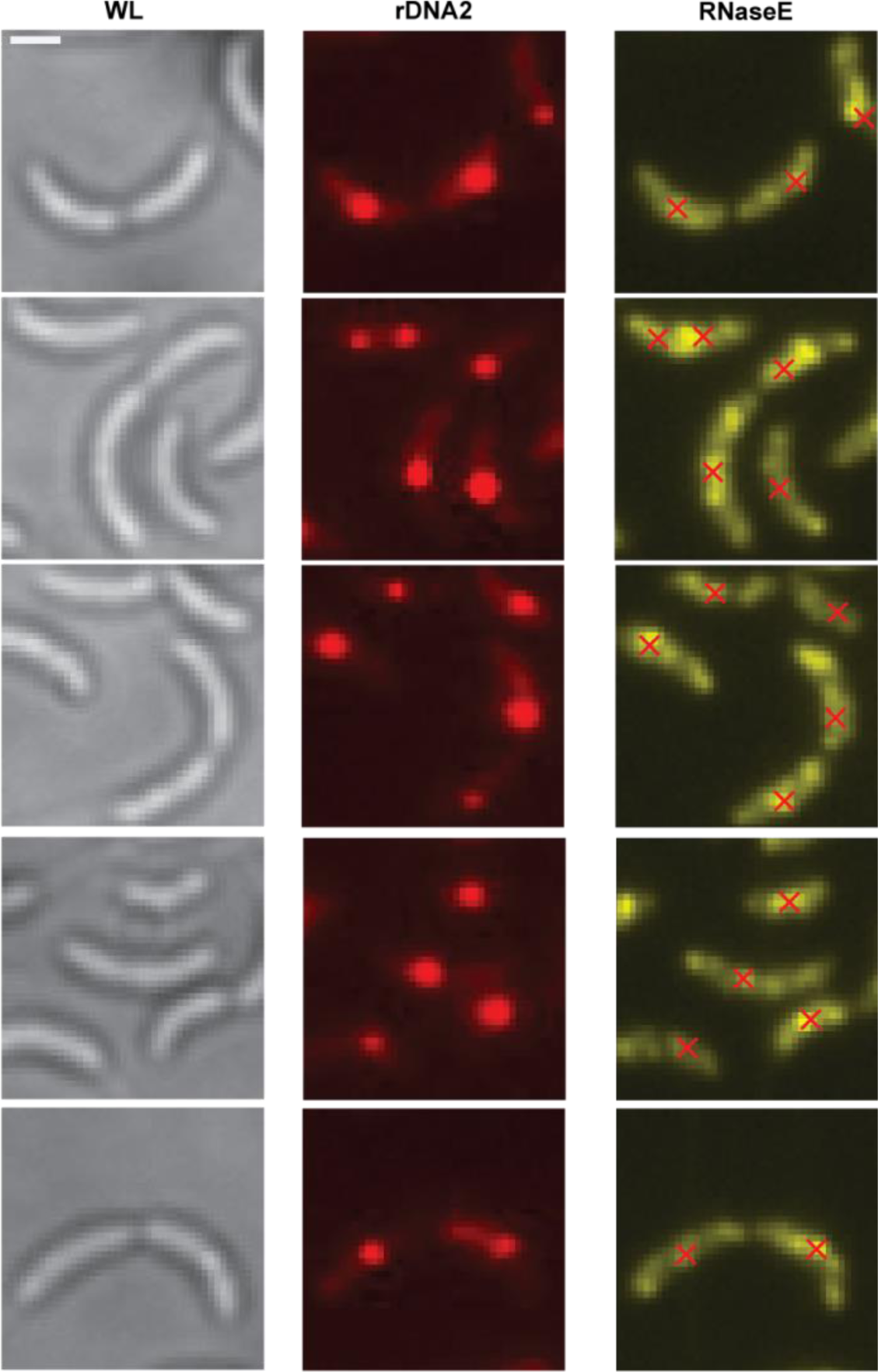
Additional DL examples of rDNA2 and RNase E colocalization in *Caulobacter*. Left column shows WL images of *Caulobacter* expressing RNase E (visualized in right column) and rDNA2-tetO/tetR (visualized in middle column). Crosses in right column show the locations of chromosomal loci displayed in middle column.

**Figure S15:**
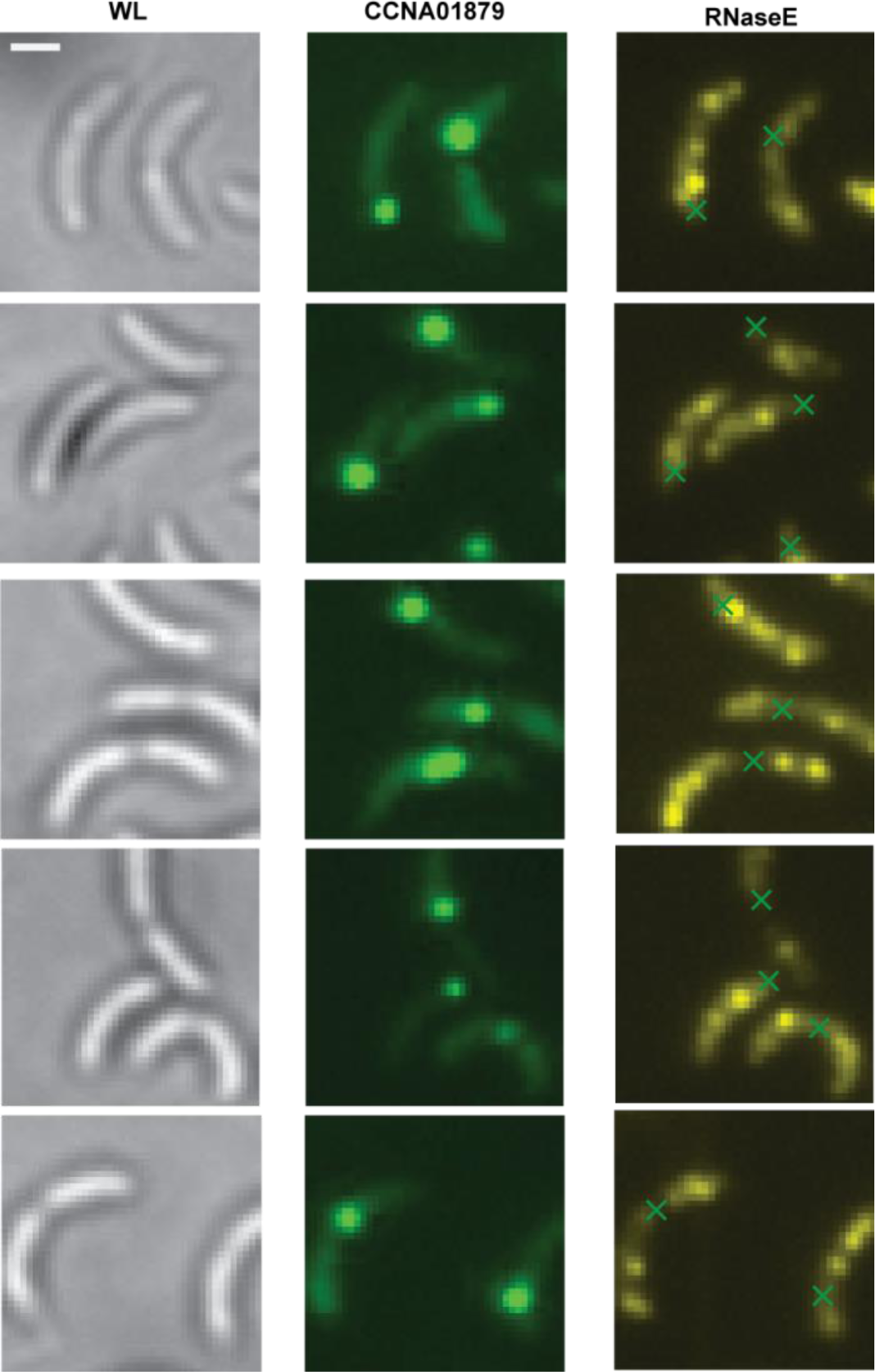
Additional DL examples of CCNA01879 and RNase E colocalization in *Caulobacter*. Left column shows WL images of *Caulobacter* expressing RNase E (visualized in right column) and CCNA01879-tetO/tetR (visualized in middle column). Crosses in right column show the locations of chromosomal loci displayed in middle column.

**Supplementary video 1**: SR reconstruction of fixed WT *Caulobacter* cell expressing RNase E-eYFP.

**Supplementary video 2**: SR reconstruction of fixed rif-treated *Caulobacter* cell expressing RNase E-eYFP.

**Supplementary video 3**: SR reconstruction of fixed WT *Caulobacter* cell expressing ribosomal protein L1-eYFP.

**Supplementary video 4**: SR reconstruction of fixed rif-treated *Caulobacter* cell expressing ribosomal protein L1-eYFP.

